# Gene editing and super-resolution microscopy reveal multiple distinct roles for ARF GTPases in cellular membrane organization

**DOI:** 10.1101/2022.05.31.494106

**Authors:** Luis Wong-Dilworth, Carmen Rodilla-Ramirez, Eleanor Fox, Steffen D. Restel, Alexander Stockhammer, Petia Adarska, Francesca Bottanelli

## Abstract

ADP-ribosylation factor (ARF) GTPases are major regulators of cellular membrane homeostasis. High sequence similarity and multiple, possible redundant functions of the five human ARFs make investigating their function a challenging task. To shed light on the roles of the different Golgi-localized ARF members in membrane trafficking, we generated CRISPR-Cas9 knock ins (KIs) of type I (ARF1 and ARF3) and type II ARFs (ARF4 and ARF5) and mapped their nanoscale localization with stimulated emission depletion (STED) super-resolution microscopy. We find ARF1, ARF4 and ARF5 on segregated nano-domains on the *cis*-Golgi and ER-Golgi intermediate compartments (ERGIC), revealing distinct roles in COPI recruitment on early secretory membranes. Interestingly, ARF4 and ARF5 define Golgi-tethered ERGIC elements decorated by COPI and devoid of ARF1. Differential localization of ARF1 and ARF4 on distal ERGICs suggests the presence of functionally different classes of intermediate compartments that could regulate bi-directional transport between the ER and the Golgi. Furthermore, ARF1 and ARF3 localize to segregated nano-domains on the *trans*-Golgi network (TGN) and TGN-derived post-Golgi tubules, strengthening the idea of distinct roles in post-Golgi sorting. This work provides the first map of the nanoscale organization of human ARF GTPases on cellular membranes and sets the stage to dissect their numerous cellular roles.

## Introduction

The regulation of membrane cellular homeostasis is essential for cell survival. Receptors and important signaling molecules rely on the secretory pathway for effective transport to their correct intracellular destination. ARF GTPases have emerged as key players in membrane homeostasis and in the regulation of global intracellular trafficking (Adarska et al., 2021). ARFs are a very ubiquitous and cross-functional family of small GTPases implicated in many cellular processes from membrane trafficking to metabolism and organization of the cytoskeleton (Jackson and Bouvet, 2014). GDP-GTP exchange factors (GEFs) load ARFs with GTP, triggering conformational changes that allow binding of downstream effectors. GTP hydrolysis mediated by GTPase activating proteins (GAPs) regulates the duration of signal output driven by ARFs (Sztul et al., 2019). Human ARFs are sub-divided in type I (ARF1 and ARF3), type II (ARF4 and ARF5) and type III (ARF6) according to sequence similarity. Type I and type II ARFs share 96% and 90% of their amino acids sequence, respectively, with 80% sequence homology shared between members of the two types. Although different ARFs bind different subsets of downstream effectors, the switch I and II regions in ARFs, which are responsible for effector binding, are almost identical (Goldberg, 1998; Pasqualato et al., 2001). ARFs high sequence similarity has limited the possibility to discern endogenous ARFs in living cells due to the lack of available antibodies, hindering our full understanding of the signaling pathways elicited by the different family members.

ARF1 is the best-known member and its role in the formation of COPI vesicles at the Golgi has been extensively studied using *in vitro* reconstitution experiments and budding assays from purified Golgi or from Golgi membranes in semi-permeabilized cells (Adolf et al., 2013; Orci et al., 1986; Reinhard et al., 2003). These experiments have led to a great molecular understanding of the ARFs on/off cycle. However, ARFs functional specificity is driven by activation on a specific membrane and recruitment of numerous effectors, requiring experiments in living cells to fully understand the cellular pathways involved. For example, ARF1, ARF4 and ARF5 have all been shown to support the formation of COPI vesicles *in vitro* (Popoff et al., 2011). However, it is unclear what happens in the cellular context and why many ARF isoforms are required. In the last two decades, ARFs have been implicated in different cellular functions beyond their role in Golgi trafficking, for which they were initially discovered. Early studies employing small interfering RNA (siRNA) to deplete ARFs in order to investigate their function, showed that type I and type II ARFs have various roles besides Golgi trafficking (Volpicelli-Daley et al., 2005). Type I and II ARFs have all been shown to have a role in TGN export and trafficking in downstream recycling endosomal compartments (Bottanelli et al., 2017; Kondo et al., 2012; Lowery et al., 2013; Manolea et al., 2010; Nakai et al., 2013; Tan et al., 2020). While ARF4 and ARF5 are associated with the ERGIC (Chun et al., 2008), they may additionally have a role in secretion and TGN-export in specialized cell types (Deretic et al., 2005; Sadakata et al., 2010).

With multiple and possible redundant functions, studying the role and endogenous distribution of ARFs in living cells has proved to be a challenging task. The observation that ARF pairs needed to be depleted from cells to yield a trafficking defect (Volpicelli-Daley et al., 2005) lead to the hypothesis that ARFs may act as heterodimers, rather than acting redundantly. A recent publication showed that GEF-dependent activation of two closely spaced ARF molecules promotes vesicle formation (Brumm et al., 2020). Additionally, ARF dimerization was shown to have a role in vesicle scission (Beck et al., 2011; Beck et al., 2008). Exogenous expression of tagged ARFs, perturbations via GTP-locked mutants, selective siRNA-mediated knock downs as well as knock outs (KO) harnessing CRISPR-Cas9 technology (Pennauer et al., 2022) have been the methods utilized so far to reveal specific cellular function of ARFs. Furthermore, the observation of the intracellular distribution of ARFs has been limited to diffraction-limited microscopy techniques. The Golgi and its neighboring organelles, such as the ERGIC, are tightly packed in the perinuclear area, making super-resolution microscopy essential for investigation of the nanoscale localization of ARFs on different nano-domains or segregating cisternae of the Golgi.

To overcome these limitations, we have generated CRISPR-Cas9 knock in (KI) cell lines of type I (ARF1 and ARF3) and type II ARFs (ARF4 and ARF5) with either the self-labelling enzymes Halo and SNAP for live-cell STED microscopy experiments or with ALFA (Gotzke et al., 2019) and V5 epitope tags to perform STED microscopy experiments in fixed cells. STED microscopy has allowed us to reveal the nanoscale distribution of each ARF GTPase and investigate their intra-Golgi distribution as well as their localization to closely juxtaposed organelles such as the *cis*-Golgi and ERGICs.

## Results

### Endogenous tagging of ARF GTPases to investigate their intracellular distribution

To be able to investigate the nanoscale distribution of endogenous ARF GTPases we generated CRISPR-Cas9 KIs of ARF1, ARF3, ARF4 and ARF5. The only type III ARF member ARF6, which has a role in endocytosis (Donaldson, 2003), was not included as this study focuses on Golgi-localized type I and II ARFs. The self-labelling enzyme tag Halo or the small epitope tag ALFA were introduced at the C-terminus of ARFs to be able to perform STED microscopy experiments.

All endogenously tagged ARFs localized to the Golgi apparatus and to vesicular and vesicular-tubular structures distributed throughout the cytoplasm (Figure 1). Halo tagging and live-cell microscopy (Figure 1a,c,e,g) are necessary to visualize ARFs tubular-vesicular structures that are otherwise disrupted upon fixation (Figure 1b,d,f,h). Complementing the live-cell microscopy performed on Halo tagged ARFs, tagging with 2xALFA allowed significant amplification of the signal of lower abundance endogenous ARFs (like ARF5) and so investigation of their intra-Golgi localization at higher resolution. Both ARF1^EN^-Halo and ARF4^EN^-Halo (Figure1a,e) highlight tubular-vesicular structures, similarly to previous observations for ARF1 (Bottanelli et al., 2017). Endogenous ARF3^EN^-Halo with a glycine/serine (GS) linker between the C-terminus of the protein and the tag displayed strong cytoplasmic localization (Supplementary figure 1). However, the addition of a long flexible linker in the form of a localization and affinity purification (LAP) tag, with eGFP substituted for the Halo tag (Cheeseman and Desai, 2005), revealed the localization of endogenous ARF3 at the Golgi and on vesicular-tubular structures (Figure 1c). Interestingly, the presence of either a GS linker or a LAP tag did not affect the localization of other endogenously tagged ARFs (Supplementary figure 1). ARF5^EN^-Halo is also enriched in the Golgi area as well as on vesicles but less prominently on tubular structures (Figure 1g). We quantified the relative abundance of all ARFs in HeLa cells and show that the abundance of ARF1 and ARF4 is similarly high, while ARF3 and ARF5 display ∼70% and ∼20% abundance compared to ARF1 and ARF4 (Figure 1j). To ensure the functionality of the fusion proteins, we generated KIs of ARFs in the haploid cell line HAP1 to show that cells do not have any discernable growth defects and show no impairment in the recruitment of the COPI and clathrin coat (Supplementary figure 2). With these KIs in hand, we set out to investigate the endogenous localization of type I and II ARFs in living and fixed cells.

**Figure 1.**
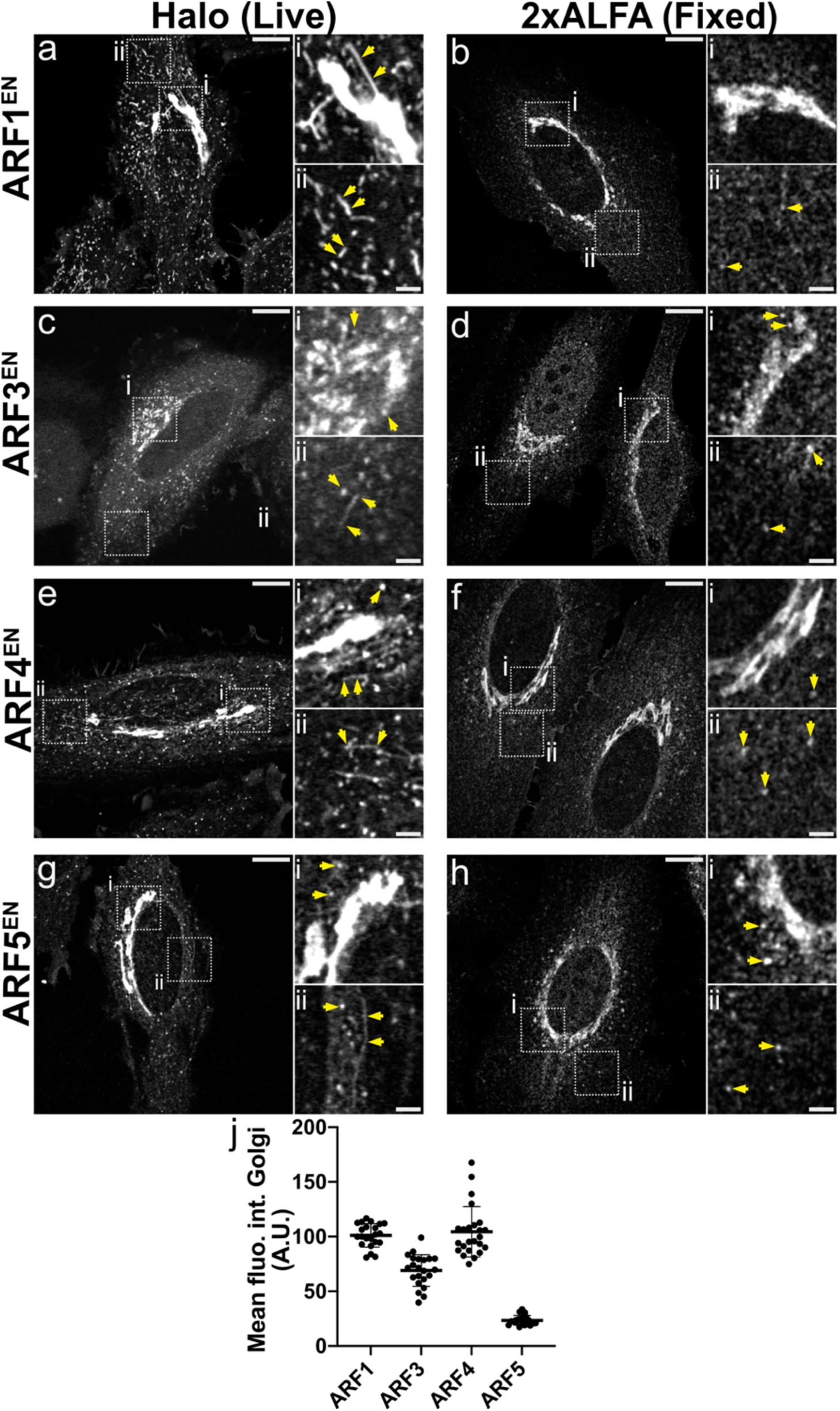
Gene editing with CRISPR-Cas9 highlights the endogenous localization of ARF GTPases. ARFs were tagged at their endogenous locus with the self-labelling enzyme Halo (a,c,e,g) or 2xALFA tag (b,d,f,h). An additional LAP tag linker was added in the case of ARF3 (c). Live cells were stained with the Halo substrate JF646-CA (a, e, g) and JFX650-CA (c). Fixed cells were immunolabeled with an anti-ALFA primary antibody and ATTO647N-conjugated secondaries. Scatter dot plots with mean and standard deviation (SD) represent the quantification of the mean fluorescence intensity at the Golgi measured in the ARF^EN^-2xALFA KI cells as described in the methods. ARF1 n=22; ARF3 n=22; ARF4 n=24; ARF5 n=20. All images were smoothed with a gaussian filter as described in the methods. CA=chloroalkane. Scale bars are 10 µm and 2 µm in the cropped images.

### A nanoscale view of the Golgi apparatus with STED super-resolution microscopy

To ultimately pinpoint the localization of ARFs, it was necessary to first obtain a map of the nanoscale localization of known Golgi and vesicular markers with STED microscopy. These included known stacking factors (GM130, Golgin97 and GRASP65) as well as the vesicular markers COPI and clathrin. The stacking factors localize to the flat part of the Golgi cisternae, which is tethered to adjacent cisternae, whereas the vesicular markers associate with the highly curved rim of the cisternae where transport intermediates are known to form and bud. STED microscopy allows easy distinction of *cis* and TGN cisternae, labelled by GM130 and Golgin97, respectively (Figure 2a). COPI clusters are observed at the outermost rim of the *cis*-Golgi cisternae labelled by GM130 (white arrows in figure 2b). STED microscopy also revealed fenestrations in the Golgi cisternae (yellow arrow in figure 2b). COPI clusters appear further away from TGN Golgin97-positive membranes in comparison to the *cis*-Golgi (Figure 2c). COPI-positive clusters represent either forming or fully formed COPI vesicles at the outmost rim of the Golgi. The stacking factors GRASP65 and GM130 localize together on the flat/stacked cisternal membranes (Figure 2d). Moreover, STED microscopy allows distinction of Golgi-associated ERGIC elements that appear to be tightly tethered to the *cis*-Golgi cisterna (white arrows in figure 2e). Interestingly, all distal (disperse throughout the cytoplasm and away from the perinuclear area) and Golgi-associated ERGICs are decorated by COPI clusters (Figure 2f). Clathrin clusters are observed in close proximity to the TGN (Figure 2g) but further apart from the *cis*-Golgi cisternae (Figure 2h). The same experiments were performed in nocodazole treated cells where the Golgi ribbon is broken down into smaller Golgi units known as ministacks (Supplementary figure 3). This approach strongly simplifies the imaging due to the more favorable geometry, allowing quantification of the distances between cisternae in downstream experiments. The observations made in intact Golgi ribbons are confirmed in nocodazole treated cells with COPI decorating the rim of *cis*-Golgi and ERGIC ministacks. We show that STED microscopy is instrumental to separate Golgi cisternae and closely juxtaposed membranes like Golgi-associated ERGICs. We could also easily distinguish nano-domains of clathrin and COPI machinery defining the curved rim of the cisternae where budding of transport intermediates occurs (Figure 2i).

**Figure 2.**
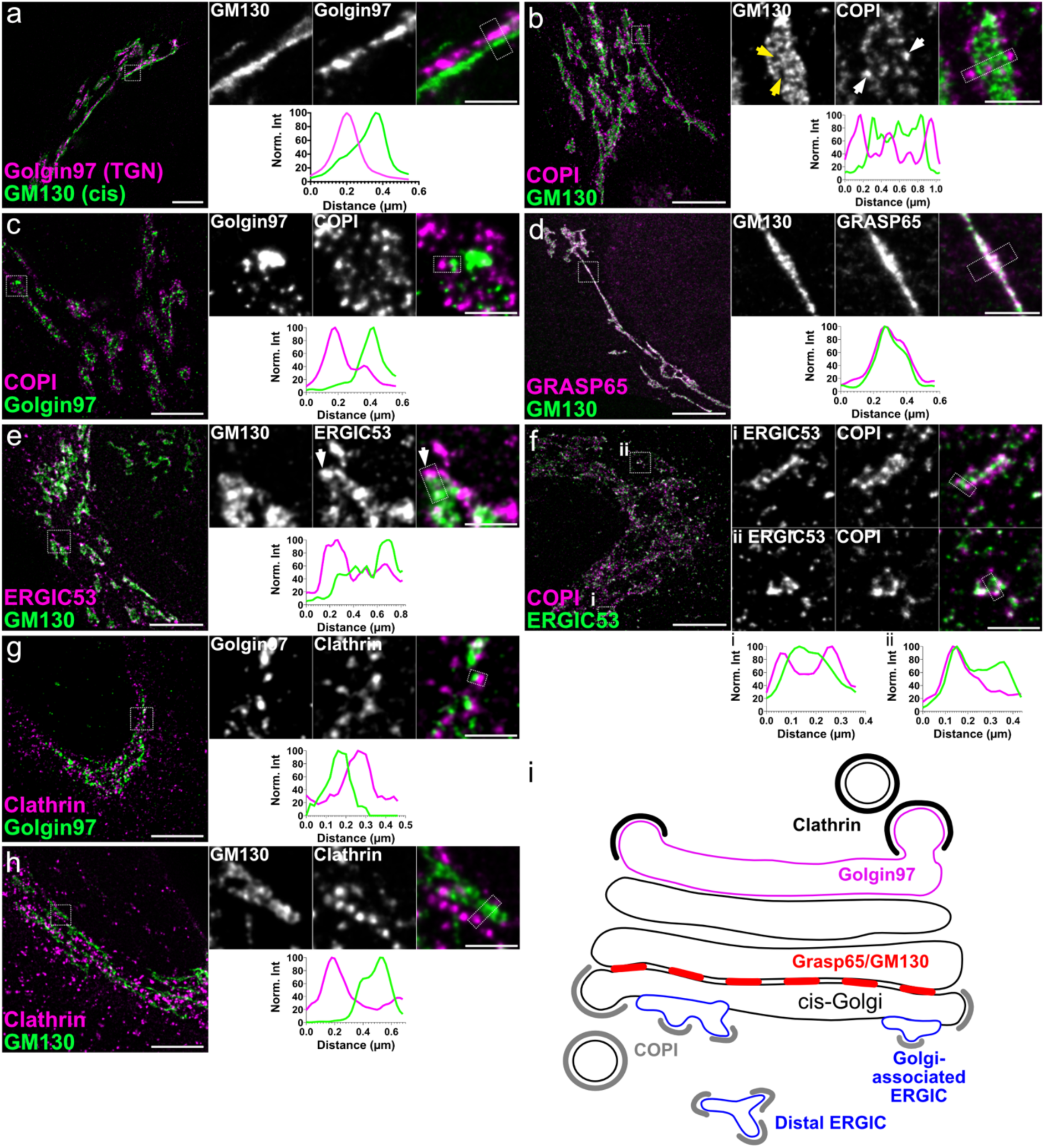
STED reveals the super-resolution localization of known Golgi and vesicular markers. HeLa cells were immunostained with antibodies against the Golgi markers indicated in the figure and secondaries labelled with either ATTO647N or AlexaFluor594 to be able to perform dual-color STED experiments. Line profiles in each panel correspond to the dotted boxes in the cropped images. All images were deconvolved and smoothed as described in the methods. Scale bars are 5 µm and 1 µm in the cropped images.

### ARF1 localizes to the *cis*-Golgi while ARF4 and ARF5 localize to Golgi-associated ERGICs

ARF1, ARF4 and ARF5 have all been localized to the ERGIC and *cis*-Golgi membranes (Chun et al., 2008) and were shown to support COPI vesicle formation in *in vitro* experiments (Popoff et al., 2011). So why are so many ARFs needed to recruit the COPI coat on early secretory membranes in living cells? COPI has a role in various trafficking steps including Golgi-to-ER transport and retrograde transport within Golgi cisternae. Additionally, a possible role for COPI in anterograde transport has been suggested and COPI is associated with Golgi-directed anterograde carriers (Stephens et al., 2000; Weigel et al., 2021). We hypothesize that different ARFs may be recruited to different nanodomains on ERGIC and *cis*-Golgi membranes to mediate different COPI-dependent trafficking steps.

To test this hypothesis, we mapped ARFs localization on the *cis*-Golgi and ERGICs with STED microscopy in fixed cells. ARF1 is associated with the *cis*-Golgi cisternae labelled by GM130 (Figure 3b,g), but excluded from *cis*-Golgi-associated ERGICs (Figure 3a,g). ARF3, the other type I ARF, is never observed on *cis*-Golgi or ERGIC membranes (Supplementary figure 4). ARF4 and ARF5 decorate the membranes of Golgi-associated and distal ERGICs (Figure 3c,e,h,i). Of the two type II ARFs, ARF4 shows some enrichment on *cis*-Golgi membranes, albeit at much lower levels than ARF1 (Figure 3d,h). Taking advantage of the easier geometry of ministacks in cells that were treated with nocodazole, we were able to quantify the distance between cisternal membranes labelled by the various ARFs, the *cis*-Golgi marker GM130 and the ERGIC marker ERGIC53 (Figure 3g-j). While ARF1 localizes to the same cisternae defined by GM130, the average ARF4 and ARF5 signal is mapped ∼400 nm away from the *cis*-Golgi, confirming their segregated localization to different cisternae (Figure 3j). The results are in agreement with observations in intact Golgi ribbons, validating ministacks as a system that greatly simplifies quantification (Tie et al., 2018; Tie et al., 2016).

**Figure 3.**
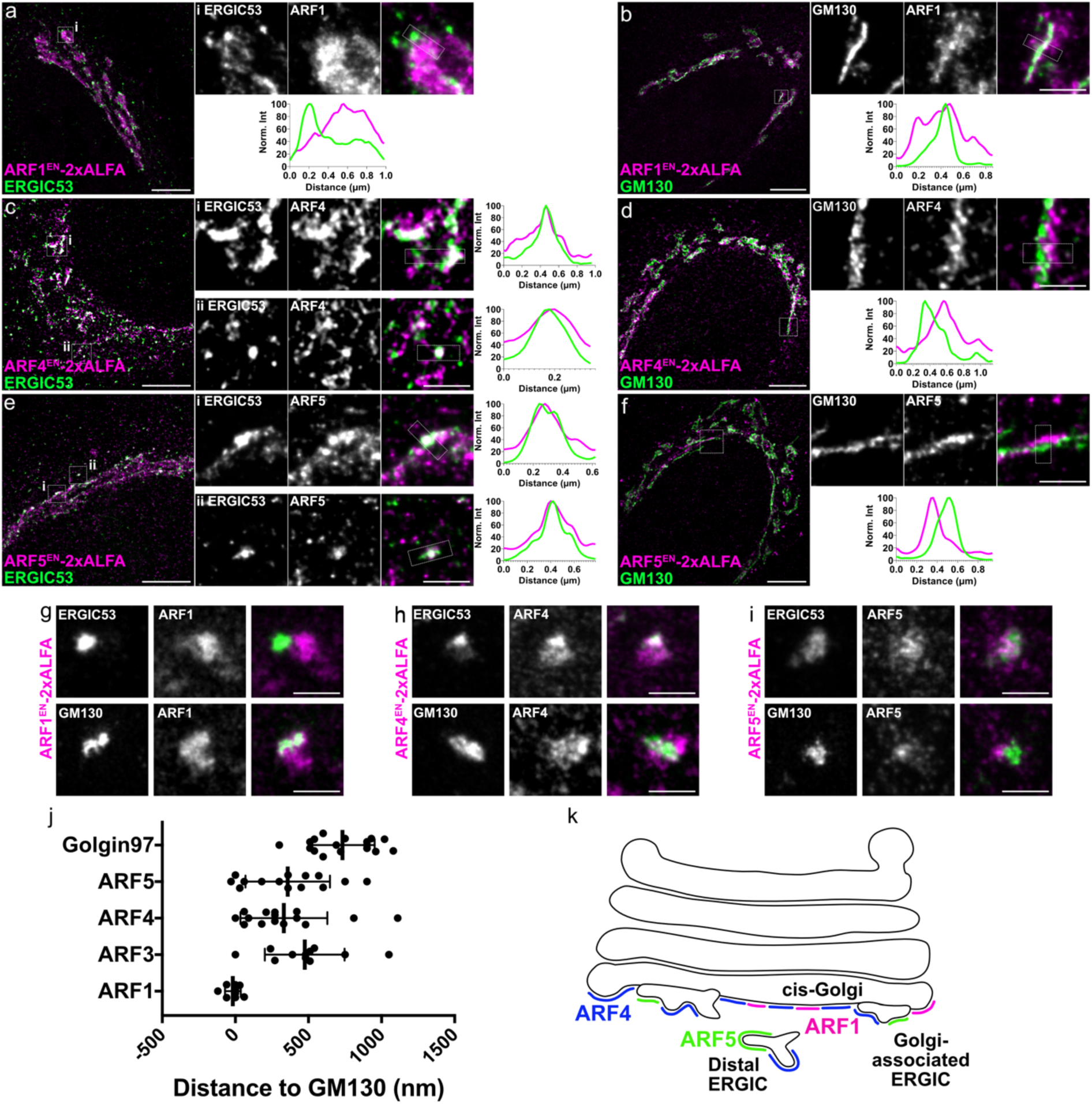
ARFs segregate on different early secretory membranes. ARF1^EN^-, ARF4^EN^-and ARF5^EN^-2xALFA KI HeLa cells were immunostained with anti-ALFA (magenta) and either anti-ERGIC53 (a,c,e, green) or anti-GM130 (b,d,f, green). Line profiles in each panel correspond to the dotted boxes in the cropped images. ARF1^EN^-, ARF4^EN^-and ARF5^EN^-2xALFA KI HeLa cells were treated with nocodazole (2.5 µg/mL) for 3h and immunostained as above. Scatter dot plot with mean and SD represents the quantification of the distance from the edge of the ARF-labelled cisternae and *cis*/ERGIC cisternae measured in the nocodazole-treated fixed cells as described in the methods. ARF1 n=9; ARF3 n=11; ARF4 n=15; ARF5 n=14; Golgin97 n=17 (j). ARF1 (and to a smaller extent ARF4) define the *cis*-Golgi membranes while ARF1, ARF4 and ARF5 are all observed on distal structures positive for ERGIC53 (k). Images were deconvolved (a-f) and smoothed (a-i) as described in the methods. Scale bars are 5 µm and 1 µm in the cropped images.

When looking at the relative distribution of ARF1, ARF4 and ARF5 in double KI cell lines, ARF1 and ARF4 are partially co-localizing on *cis*-Golgi membranes while distinct ARF4-positive Golgi-associated ERGICs devoid of ARF1 are clearly visible (Supplementary figure 5a). Additionally, ARF4 and ARF5 exclusively localize to segregated nanodomains on the ERGIC (Supplementary figure 5b), suggesting a functional differentiation between the two type II ARFs. Altogether, this data suggests that ARF1, ARF4 and ARF5 segregate on different early secretory membranes (Figure 3k).

### Live-cell STED highlights segregation of ARF1 and ARF4 on different sub-classes of tubular-vesicular early secretory elements

As tubular-vesicular elements at the Golgi and in the cytoplasm are disrupted upon fixation (Figure 1), we performed live-cell STED experiments to better characterize the distribution of ARFs on Golgi-associated and distal tubular-vesicular ERGIC and *cis*-Golgi-derived structures. For dual-color live-cell STED experiments, Halo and SNAP tag fusions are labelled with cell permeable Halo chloroalkane (CA) and SNAP benzylguanine (BG) substrates conjugated to STED-compatible organic dyes (Bottanelli et al., 2016; Stockhammer and Bottanelli, 2021). Multi-color live-cell STED microscopy of gene-edited βCOP^EN^-Halo and transiently expressed SNAP-ERGIC53 highlights COPI clusters on tubular-vesicular ERGIC elements associated with the Golgi ribbon (Supplementary figure 6i) and with distal-ERGIC structures further away from the peri-nuclear area (Supplementary figure 6ii, iii). Golgi-associated ERGIC elements are devoid of ARF1^EN^-Halo and appear as tubular-vesicular structures or cisternae-like structures tethered to the *cis*-Golgi cisterna labelled by ARF1 (white arrows in Figure 4a). In addition to defining the *cis*-Golgi cisternae, ARF1^EN^-Halo also localizes to a fraction of distal ERGICs (Figure 4a-iii) which were not easily detectable in fixed cell imaging. ARF4^EN^-Halo decorates Golgi-associated ERGICs and a larger sub-set of distal ERGIC53-positive structures (Figure 4b). As both ARF4 and ARF1 localize to similar tubular-vesicular structures at the Golgi, we decided to compare the localization of ARF1 and ARF4 in a double KI cell line where ARF4 and ARF1 have been endogenously tagged with Halo and SNAP tag, respectively (Figure 4c). ARF4^EN^-Halo is present on the *cis*-Golgi cisternae, albeit at a lower level than ARF1, as seen from the partial overlap with ARF1^EN^-SNAP signal (Figure 4c-i). However, ARF4^EN^-Halo is strikingly enriched on Golgi-associated ERGICs, which are completely devoid of ARF1^EN^-SNAP (Figure 4c-i). Additionally, ARF1^EN^-SNAP and ARF4^EN^-Halo define segregated tubular vesicular elements seen in close proximity to the Golgi or in the cell periphery (Figure 4c-ii,iii). As ARF1 and ARF4 defined different populations of ERGICs, we quantified the number of distal ERGICs positive for the various ARFs in whole cell confocal micrographs (Figure 4d and Supplementary figure 7). While ∼70% of the structures are positive for ARF4, only ∼35% are positive for ARF1 and ∼35% are positive for both ARF1 and ARF4 (Figure 4d). Interestingly, when focusing on the nanoscale localization of ARF1 and ARF4 on distal ERGICs with live-cell STED, the two GTPases are seen on segregating nano-domains on the same structure (Figure 4e). These results suggest a differential sorting function for ARF1 and ARF4 at the *cis*-Golgi-ERGIC interface at the level of sorting tubular-vesicular elements. While ARF1 tubules are retrograde in nature (Bottanelli et al., 2017), ARF4 could contribute to anterograde flow or provide an early recycling platform from the ERGIC to the Golgi (Figure 4f).

**Figure 4.**
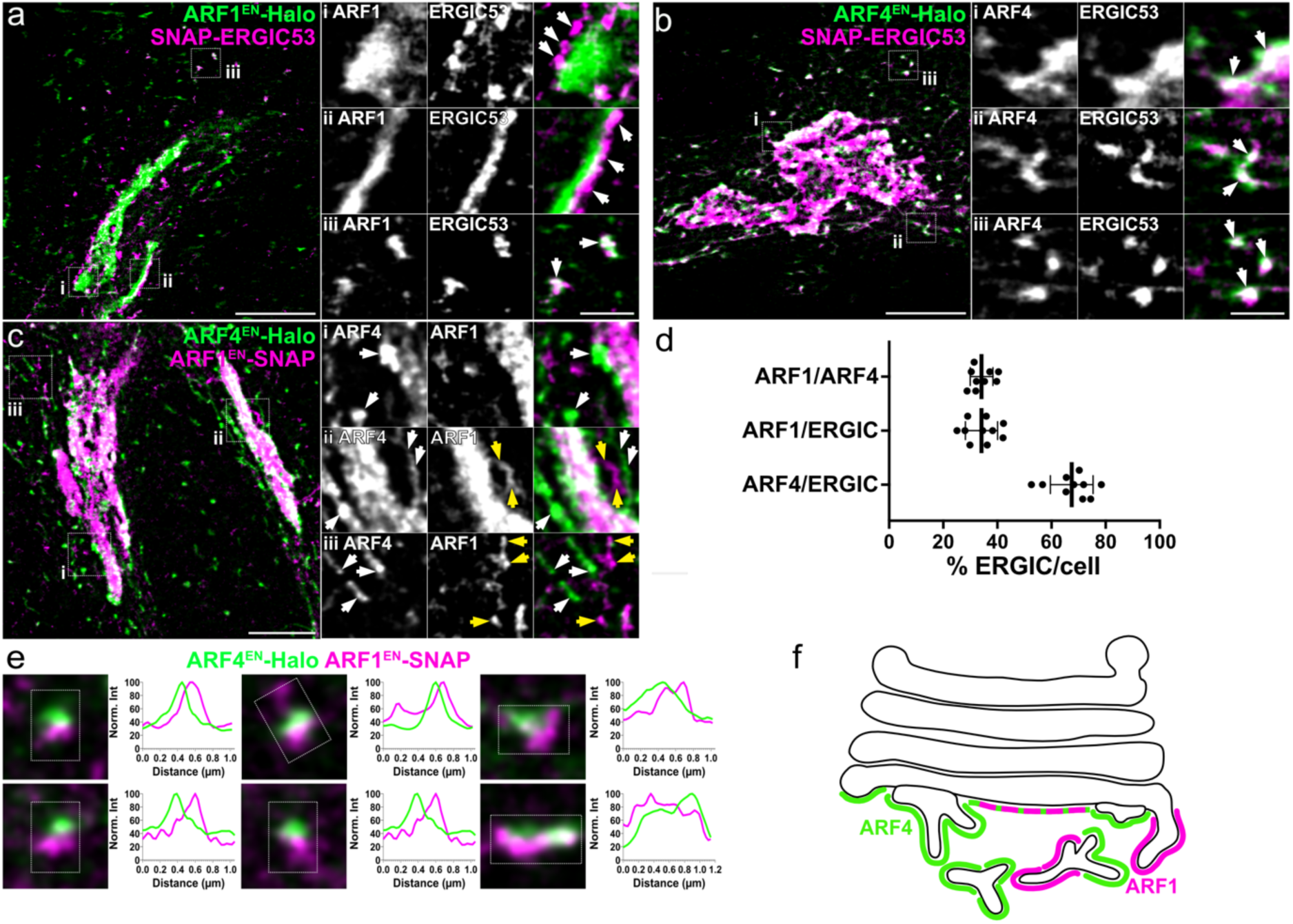
ARF1 and ARF4 define different populations of ERGICs. ARF1^EN^-Halo and ARF4^EN^-Halo KI cells were transfected with a plasmid encoding for SNAP-ERGIC53 (a, b). Live cells were stained with the Halo substrate JF571-CA and the SNAP substrate JFX650-BG (a, b). ARF1^EN^-SNAP and ARF4^EN^-Halo double KI cells were labelled with JF571-CA and JFX650-BG (c,e). The percentage of distal ERGIC positive for either ARF1 or ARF4 was calculated by counting all ARF1/ARF4 positive distal ERGICs divided by the total amount of ERGICs. The percentage of ARF1-and ARF4-positive distal structures was calculated by counting all ARF1 and ARF4 positive structures divided by the total amount of distal ARF4 structures (d). ARF4/ERGIC n=10; ARF1/ERGIC n=10; ARF1/ARF4 n=9. Error bars represent mean and SD. ARF4 defines Golgi-associated ERGICs and ARF1/ARF4 segregate on different populations of distal ERGICs and on nano-domains on the same ERGIC (f). All images were deconvolved and smoothed as described in the methods. CA=chloroalkane, BG= benzylguanine. Scale bars are 5 µm and 1 µm in the cropped images.

### Different ARF isoforms support COPI recruitment on different ERGIC/Golgi membranes

The differential localization of ARF1, ARF4 and ARF5 to the *cis*-Golgi and ERGICs indicates that different ARFs may be responsible for COPI recruitment on different early secretory membranes. While all ARFs were shown to recruit COPI *in vitro*, the localization and proximity of COPI to each ARF isoform has never been explored in living cells. Taking advantage of our gene editing approach combined with live-cell STED, we generated double KIs where ARFs were tagged with a Halo tag whereas the β subunit of the COPI coat has been tagged with a SNAP tag to investigate the localization of ARF isoforms and COPI machinery (Figure 5 and Supplementary figure 8). ARF1^EN^-Halo is seen in close proximity to COPI coated vesicular structures that define the *cis*-Golgi rims (Figure 5a-i). In line with previous work, ARF1^EN^-Halo defines tubular-vesicular non-ERGIC structures positive for COPI emanating from the Golgi (Figure 5a-ii) and in the cell periphery (Figure 5a-iii). These tubules were shown to carry cargoes like KDEL receptor and to be retrograde in nature (Bottanelli et al., 2017). However, ARF1^EN^-Halo is rarely enriched in COPI-positive structures, suggesting that upon COPI recruitment on the *cis*-Golgi, GAP-dependent GTP hydrolysis and ARF release occurs very quickly. Live-cell STED is again instrumental in showing that Golgi-associated ARF4^EN^-Halo ERGIC tubular-vesicular elements are also decorated with COPI, similar to observations for ARF1 (Figure 5b). However, ARF4^EN^-Halo is often observed enriched in COPI-positive structures (Figure 5b-ii, yellow arrows). Quantification of the average distance between COPI clusters and the membranes defined by the various ARFs in nocodazole treated cells, shows that ARF4 and ARF5 are found to be on average (∼-30 nm and 5 nm, respectively) more enriched near and within COPI clusters than ARF1 (∼100 nm), confirming the initial qualitative observations (Figure 5c-d). A differential enrichment of ARF isoforms in COPI structures in living cells hints at a possible differential regulation by GAPs at the *cis*-Golgi (ARF1) and ERGIC (ARF4 and ARF5).

**Figure 5.**
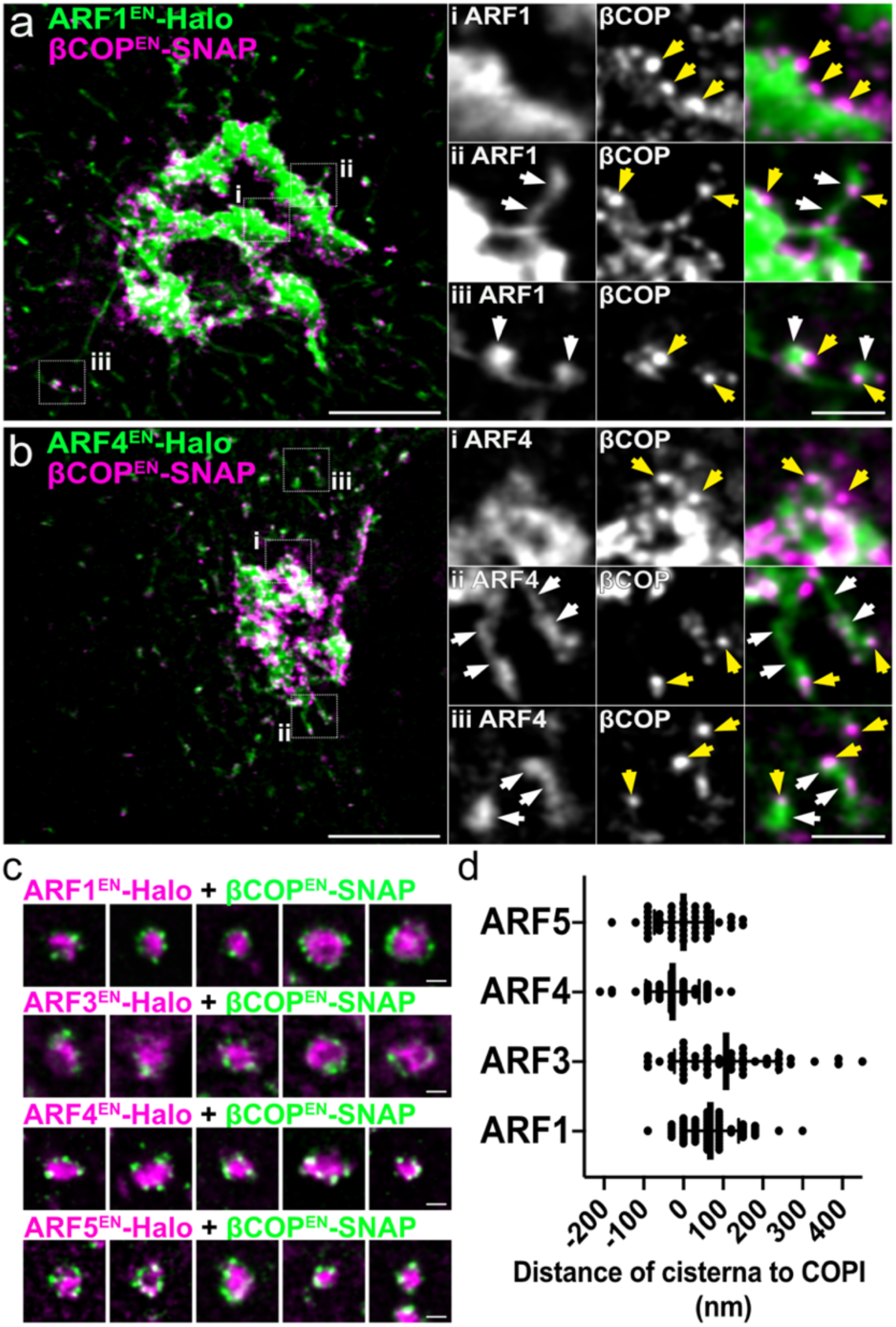
Live-cell STED reveals segregated ARF1 and ARF4 sub-populations of tubular vesicular structures defined by COPI machinery. ARF1^EN^/ARF4^EN^-Halo (green) and βCOP^EN^-SNAP (magenta) double KI cells were labelled with JF571-CA and JFX650-BG (a,b). Double KI cell lines ARF1^EN^/ARF3^EN^/ARF4^EN^/ARF5^EN^-Halo (magenta) and βCOP^EN^-SNAP (green) were labelled with JFX650-CA and JF585-BG and subsequently treated with nocodazole (2.5 µg/mL) for 3 h (c). Scatter dot plot with mean and SD represents the quantification of the distance from the edge of the ARF-labelled cisternae to the center of COPI vesicles measured in nocodazole-treated live cells as described in the methods (d). ARF1 n=62; ARF3 n=49; ARF4 n=49; ARF5 n=48. All images were deconvolved and smoothed as described in the methods. CA=chloroalkane, BG= benzylguanine. Scale bars are 5 µm and 1 µm in the cropped images.

### Type II ARFs ARF1 and ARF3 are the sole ARFs localizing to TGN membranes

After establishing the role of different ARFs in sorting at early secretory organelles, we next wanted to test their role in export from the TGN. As all ARFs have been implicated in post-Golgi trafficking (Adarska et al., 2021) we set out to understand which ones localize to the most distal TGN cisternae and to investigate their spatial relationship with the clathrin machinery. Firstly, the localization of the different ARF members endogenously tagged with 2xALFA was tested with the TGN marker Golgin97. To our surprise, only type I ARFs, ARF1 and ARF3, are seen on TGN cisternae (Figure 6a-d). Taking advantage of nocodazole-treated ministacks and their easier geometry, we show that ARF3 is exclusively enriched around TGN membranes, while ARF1 signal is more disperse, confirming its widespread Golgi localization from *cis* to *trans* (Figure 6e-f). Interestingly, cisternae positive for type II ARFs (ARF4 and ARF5) are on average ∼700 nm away from TGN membranes defined by Golgin97, similar to the distance observed between GM130 and the TGN. While in HeLa cells ARF4 and ARF5 do not associate with the TGN, cell-type dependent roles in TGN-export may be present, as previously reported (Deretic et al., 2005; Sadakata et al., 2010).

**Figure 6.**
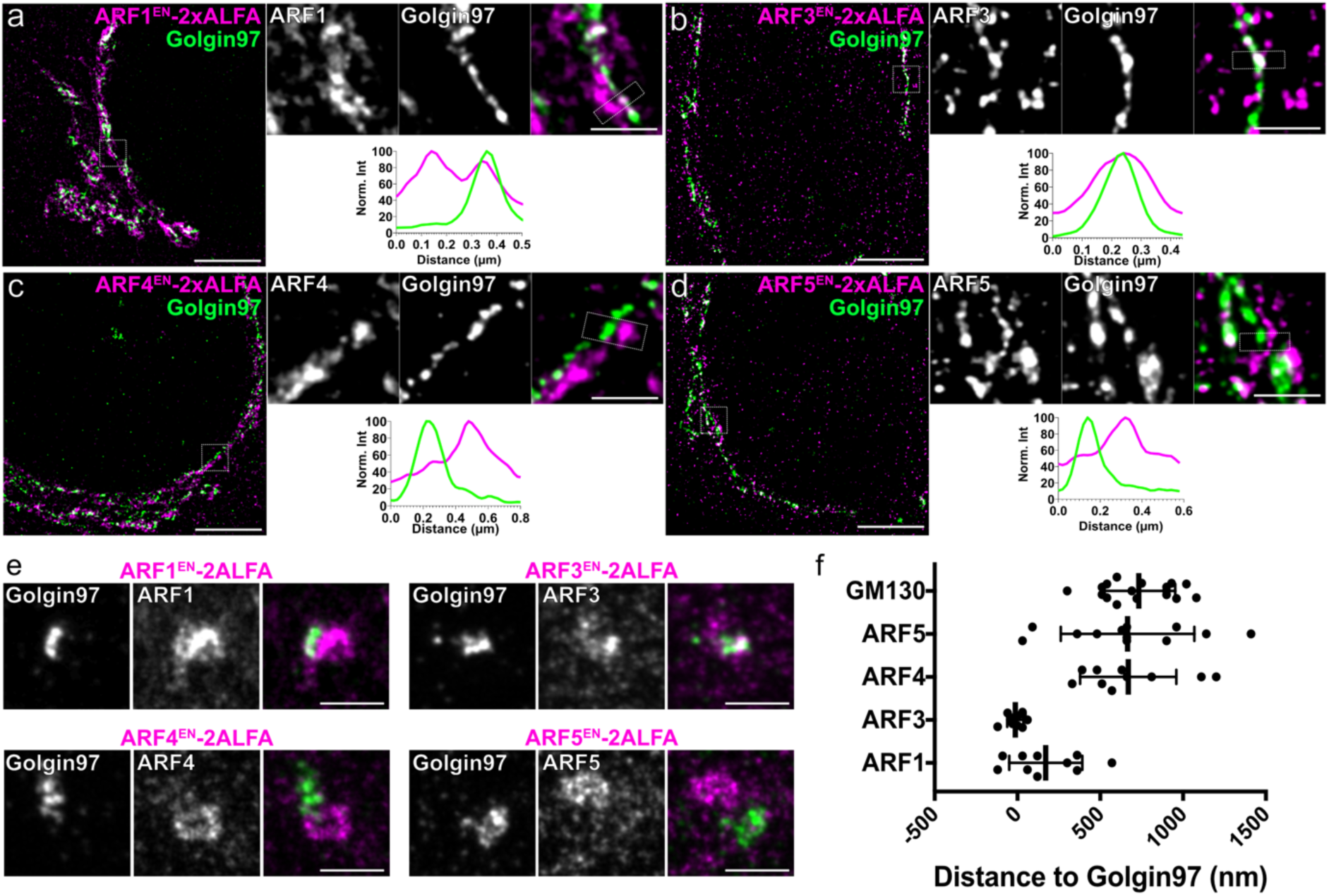
ARF1 and ARF3 are the sole TGN-localized ARF members. ARF1^EN^-, ARF3^EN^-, ARF4^EN^-and ARF5^EN^-2xALFA KI HeLa cells were immunostained with anti-ALFA tag (a-e; magenta) and anti-Golgin97 (a-e; green). ARF1^EN^-, ARF3^EN^-, ARF4^EN^-and ARF5^EN^-2xALFA KI HeLa cells were treated with nocodazole (2.5 µg/mL) for 3h and immunostained as above (e). Scatter dot plot with mean and SD represents the quantification of the distance from the edge of the ARF-labelled cisternae and Golgin97-labelled TGN cisternae measured in nocodazole-treated fixed cells as described in the methods (f). ARF1 n=10; ARF3 n=12; ARF4 n=10; ARF5 n=12; GM130 n=17. Images were deconvolved (a-d) and smoothed with a gaussian filter (a-e) as described in the methods. Scale bars are 5 µm and 1 µm in the cropped images.

### Live-cell STED shows that ARF1 and ARF3 define segregating nano-domains on the TGN and TGN-derived clathrin positive tubular vesicular carriers

An important function of ARFs is the recruitment of clathrin and adaptors to the TGN for enrichment of cargoes and formation of post-Golgi carriers. Transport intermediates both vesicular and tubular in nature have been reported (Stalder and Gershlick, 2020). We previously showed that some anterograde cargoes leave the TGN in ARF1-positive tubular vesicular carriers (Bottanelli et al., 2017). Taking advantage of live-cell STED to preserve sensitive tubular structures we wanted to test whether any other ARF define TGN-associated tubular vesicular elements defined by the clathrin coat. To this end, we generated double KIs of all ARFs (Halo tagged) and clathrin light chain (SNAP tagged, Figure 7). As previously shown, ARF1^EN^-Halo defines TGN-derived tubular vesicular structures decorated by clathrin (Figure 7a). Like ARF1, clathrin clusters are seen adjacent to cisternae defined by highly homologous ARF3^EN^-LAP-Halo (Figure 7b-i). ARF3^EN^-LAP-Halo also defines tubular-vesicular structures decorated by clathrin (Figure 7b-ii-iii). Clathrin clusters (associated with the Golgi, endosomes or the plasma membrane) densely populate the cell cytoplasm, making it hard to define the proximity of ARFs to clathrin in cells with intact Golgi ribbons. To overcome this issue, we again took advantage of nocodazole-induced ministacks and their simpler imaging geometry (Figure 7e). Quantification of the distance of clathrin clusters from the edge of the cisternae defined by the various ARFs reveals that ARF4 and ARF5 are the most distal (∼200 nm on average) ARFs from clathrin-positive structures (Figure 7e-f). ARF1 is on average ∼100 nm away from a TGN-associated clathrin vesicle, while ARF3 is the most enriched.

**Figure 7.**
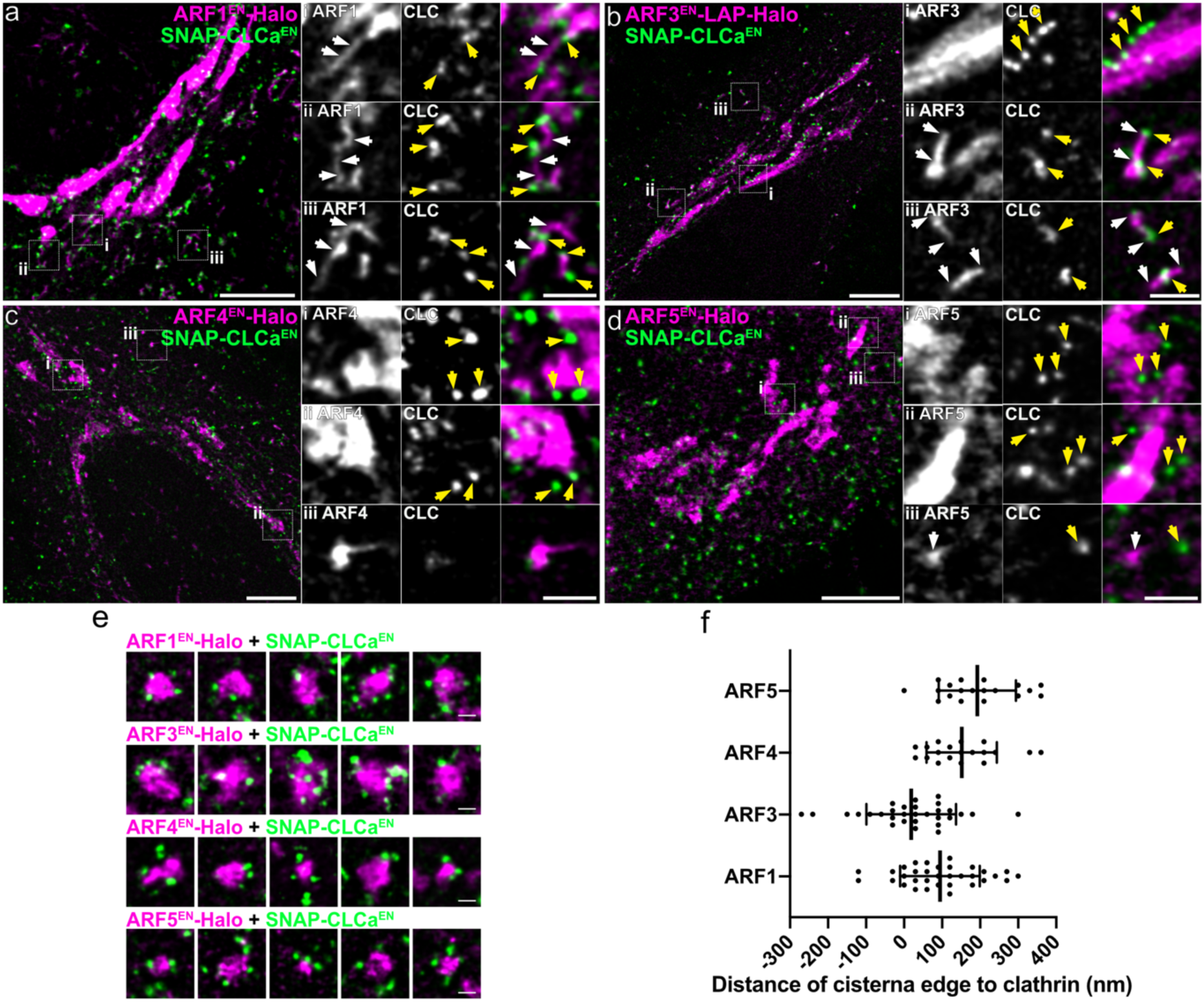
Type I ARFs define tubular-vesicular clathrin-positive structures on the TGN and the cell periphery. ARF1^EN^-, ARF3^EN^-, ARF4^EN^-and ARF5^EN^-Halo (magenta) and SNAP-CLCa^EN^ (green) double KI cells were labelled with JF571-CA together with JFX650-BG (a,c) or JFX650-CA and JF585-BG (b,d). Double KI cell lines were labelled with JFX650-CA and JF585-BG and then treated with nocodazole (2.5 µg/mL) for 3h (e). Scatter dot plot with mean and SD represents the quantification of the distance from the edge of the ARF-labelled cisternae to the center of clathrin vesicles measured in nocodazole-treated live cells as described in the methods (f). ARF1 n=33; ARF3 n=30; ARF4 n=19; ARF5 n=19. All images were deconvolved and smoothed as described in the methods. CA=chloroalkane, BG= benzylguanine. Scale bars are 5 µm and 1 µm in the cropped images.

While ARF1 and ARF3 differ only for 7 amino acids at the very N-and C-termini of the proteins, a functional differentiation for the two GTPases has been suggested (Manolea et al., 2010; Volpicelli-Daley et al., 2005). To test the extent of ARF1 and ARF3 co-localization on the TGN membranes and post-Golgi trafficking intermediates we generated double KIs where both ARF1 and ARF3 were endogenously tagged with Halo and LAP-SNAP tags, respectively. Many areas of the Golgi defined by ARF1 are devoid of ARF3, representing *cis*-Golgi cisternae (Figure 8a). Interestingly, while ARF1 localization is more evenly distributed on the cisternae, ARF3 is organized in clusters and on nano-domains completely devoid of ARF1 (Figure 8a-i). The two GTPases also segregate on TGN-derived clathrin-positive tubular trafficking intermediates emanating from the Golgi (Figure 8-ii, iii, iv). This suggests the presence of TGN-derived Golgi fragments where ARF1 and ARF3 have segregated downstream sorting functions (Figure 8b). Although ARF1 is often considered to be the master regulator for clathrin recruitment at the TGN and ARF-dependent post-Golgi sorting, our results points towards an unexplored role of ARF3 in a pathway that segregates from ARF1-mediated trafficking.

**Figure 8.**
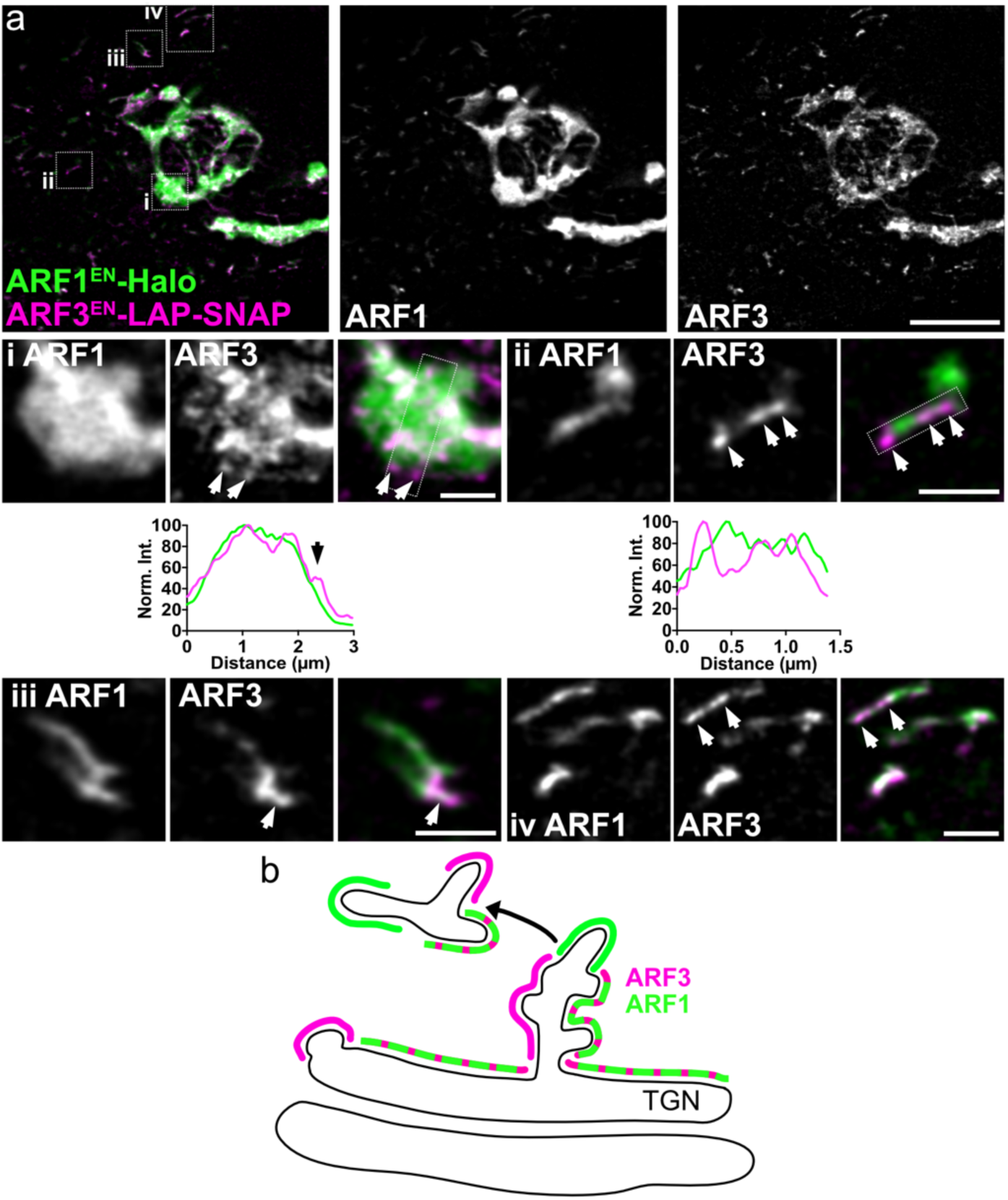
ARF1 and ARF3 segregating nano-domains on the TGN and TGN-derived tubular vesicular trafficking intermediates. ARF1^EN^-Halo and ARF3^EN^-LAP-SNAP double KI cells were labelled with JF571-CA and JFX650-BG (a). Crops highlight segregation of gene edited ARF1 and ARF3 on the Golgi (i) and on TGN-derived tubular vesicular trafficking intermediates (ii-iv). ARF3 nano-domains are highlighted by white arrows in the crops or a black arrow in the line profile graph. The localization of the two GTPases suggest that ARF1 and ARF3 have segregated sorting functions on TGN-derived trafficking intermediates (b). Images were deconvolved and smoothed with a gaussian filter as described in the methods. The brightness in the crops was enhanced to highlight the dim distal structures. CA=chloroalkane, BG= benzylguanine. Scale bars are 5 µm and 1 µm in the cropped images.

## Discussion

In this report, we demonstrate how defining the nanoscale localization of endogenous ARF GTPases has proven to be beneficial to understand their multiple cellular roles. With the help of gene editing and super-resolution microscopy we could finally map ARFs to their intracellular localization in HeLa cells (Figure 9), necessary for future understanding of how ARFs act in concert with their regulatory GAPs and GEFs to trigger specific cellular responses at specific locations inside living cells. Analysis of the localization of endogenously tagged ARFs highlights functional differences for ARF1, ARF4 and ARF5 at the ER-Golgi interface and ARF1 and ARF3 in post-Golgi trafficking. ARF1 is the only ARF with ubiquitous localization throughout the Golgi stack, which suggests it may be the sole ARF involved in intra-Golgi transport. Even higher resolution microscopy will be required to resolve transport intermediates connecting cisternae and whether they carry only retrograde or also anterograde-directed cargoes (Orci et al., 2000). Localization of ARF1 and ARF3 on partially segregating nano-domains on the TGN and post-Golgi tubular-vesicular clathrin-positive structures indicate a differential role for type I GTPases in sorting on post-Golgi tubules. We want to stress the importance of being able to perform super-resolution microscopy experiments in living cells, as many secretory elements like ERGICs and TGN-derived tubular vesicular-structures are disrupted upon fixation, hindering the investigation of trafficking mechanisms.

All type I and II ARFs were shown to support the recruitment of the COPI coat *in vitro* (Popoff et al., 2011) making us wonder why multiple ARF isoforms are needed and whether their function is redundant. Endogenous tagging combined with super-resolution STED microscopy reveals that ARF1 and type II ARF4 and ARF5 segregate on early secretory membranes (Figures 3-5). ARF1, and to a less extent ARF4, localize to the *cis*-Golgi cisternae. Interestingly, type II ARFs are enriched on ERGIC tubular-vesicular elements that appear tethered to the *cis*-Golgi cisternae, completely devoid of ARF1. ARF1, ARF4 and ARF5 also define COPI-positive distal ERGIC membranes in the cytoplasm, with a clear fraction of ERGICs which are ARF4-positive only. Interestingly, STED microscopy highlights that ARF4 and ARF1 localize to segregating nanodomains on ERGICs, whereas the two GTPases are seen colocalizing with confocal microscopy. Our data suggests that ARF1, ARF4 and ARF5 have differential trafficking roles in the early secretory pathway. While ARF1 tubular-vesicular structures forming at the *cis*-Golgi are loaded with retrograde KDEL receptor cargoes (Bottanelli et al., 2017), we speculate that ARF4-ARF5 ERGICs have a role in either anterograde ERGIC-to-Golgi transport or ERGIC-to-ER recycling. ARF1 would mediate recycling of ER residents from the *cis*-Golgi, while early retrieval from the ERGIC would be mediated by ARF4 and ARF5. This model would agree with mass spectrometry analysis showing that COPI vesicles generated with different ARFs have the same content (Adolf et al., 2019). It is unclear whether ARF/COPI are implicated in anterograde ERGIC-to-Golgi transport or simply provide an earlier retrieval platform. Interestingly, ARF4 and ARF5 localize to segregating nano-domains on ERGICs, suggesting a functional differentiation within type II ARFs. Further work would be needed to pinpoint the exact role of ARF4 and ARF5 on ERGIC membranes. Another interesting question is whether different COPI isoforms (Moelleken et al., 2007) may be recruited by different ARF isoforms via yet unknown cytosolic adaptors. Scly1 has been proposed to exclusively interact with ARF4 and ARF5 and lead to the recruitment of γ_2_-COP (Hamlin et al., 2014). Further analysis of the interactome of ARFs will be required to identify possible adaptors specific for other ARFs and tools presented here will be instrumental to test interactions and super-resolved localization of the possible adaptor-ARF pairs.

**Figure 9.**
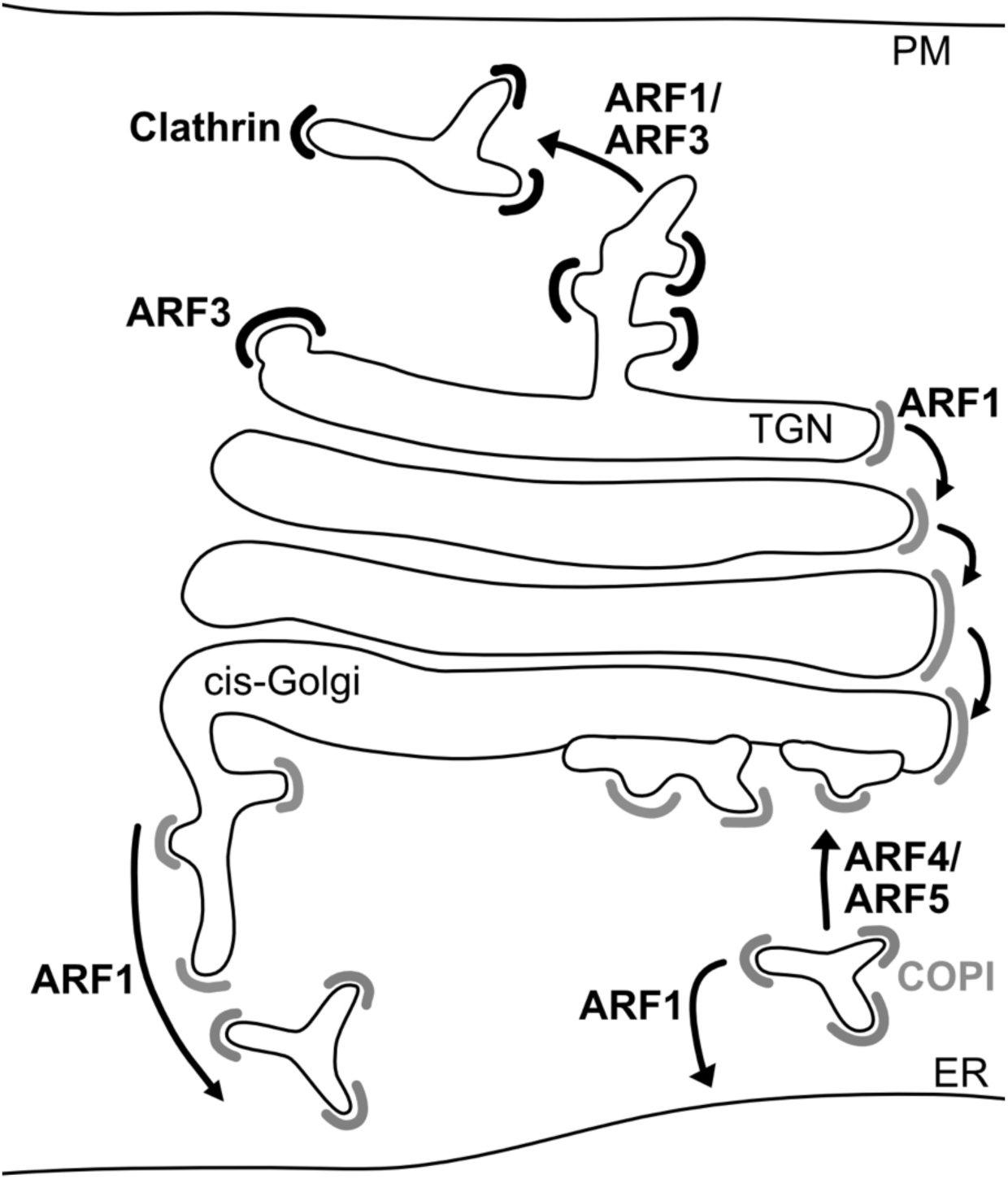
Model proposing distinct functions of different ARF isoforms in ER-Golgi transport and TGN export.

Next, we were questioning what leads to the differential recruitment of ARF1, ARF4 and ARF5 on early secretory membranes. Only a single ARF GEF is observed on the *cis*-Golgi/ERGIC membranes making it unlikely that differential recruitment is driven by the activating GEF (Monetta et al., 2007). A 16 amino acid sequence in helix α3 of ARF1 has been shown to bind to the SNARE membrin and be responsible for the recruitment of ARF1 to early Golgi cisternae (Honda et al., 2005). Additionally, p24 family proteins are known to lead to the recruitment of GDP-bound ARF1 (Gommel et al., 2001; Majoul et al., 1998) raising the possibility that different p24 proteins may bind different ARFs at different cellular locations. Structural information shows that while ARF1 and ARF6 are structurally very similar in the GTP-bound form, they differ in their GDP-bound form (Goldberg, 1998; Pasqualato et al., 2001). This suggests that the highly homologous ARF1, ARF4 and ARF5 could differentiate between a different subset of effectors in their GDP bound forms. Investigation of the proximity of COPI clusters to the various ARFs highlighted that ARF1, ARF4 and ARF5 are differentially enriched in COPI clusters (Figure 5), with ARF4 and ARF5 being the most enriched. GTP hydrolysis on ARFs and membrane dissociation could occur very quickly after COPI recruitment, as ARF GAP2 and ARF GAP3 are known to bind to the COPI coat and binding enhances GAP activity (Arakel et al., 2019; Weimer et al., 2008). Interestingly, COPI association to the membrane is maintained, despite loss of ARF in the COPI-positive clusters. So far very little is known about the dynamics of COPI vesicles uncoating *in vivo*. Our work suggests that ARF1 may dissociate from the membrane earlier than the COPI coat. Additionally, ARF4 and ARF5 are more strongly enriched in COPI clusters than ARF1 suggesting that the regulation of ARF1 at the *cis*-Golgi and ARF4/ARF5 at the ERGIC may be controlled by different ARF GAPs.

It has also been hypothesized that ARFs act as heterodimers. ARF1 dimerization has been shown to be important for fission of carriers (Beck et al., 2011; Beck et al., 2008) and dimerization of an ARF GEF in plant cells has been shown to lead to the recruitment and activation of closely positioned ARF pairs (Brumm et al., 2020). Additionally, knock down of single ARF GTPases failed to yield any phenotype, while knock down of ARF pairs affected specific trafficking steps (Kondo et al., 2012; Nakai et al., 2013; Volpicelli-Daley et al., 2005). However, CRISPR-Cas9 KO cell lines of ARF1 and ARF4 were recently shown to have an effect on Golgi membrane homeostasis and retrograde Golgi-to-ER transport, while in other double KO cell lines no additional defects were observed (Pennauer et al., 2022). Interestingly, ARF1 and ARF4 could not be deleted simultaneously. Though at a lower level, ARF4 is observed on the *cis*-Golgi membranes with ARF1 (Figure 4c) raising the possibility that ARF1 and ARF4 may act together to mediate *cis*-Golgi-to-ER transport and that they could compensate for each other when only one is depleted. ARF4 and ARF5 partially co-localize on the ERGIC membranes, as ARF1 and ARF3 overlap on the TGN. This is a hint that these ARFs may have shared as well as non-overlapping functions. However, the observed overlap may be due to initial association/activation to the same membranes, followed by segregation triggered by association with different sub-sets of effectors.

Although ARF4 and ARF5 have been implicated in post-Golgi trafficking, imaging of endogenously tagged ARFs shows that only ARF1 and ARF3 are associated to the TGN (Figure 6). A limitation of this study is that all editing and imaging is carried out in HeLa cells, in which cell-type specific functions of ARF4 and ARF5 won’t be present. For example, ARF4 has been implicated in trafficking to the cilium (Deretic et al., 2005). However, we cannot reconcile our observation with a function of ARF4 at endosomes (Nakai et al., 2013) as ARF4 exclusively associates with ERGIC membranes. Our editing tools, consist of a resistance cassette that allows for selection of successfully edited cells (Stockhammer et al., 2021) and therefore will allow to easily edit different cell types to facilitate the discovery of cell-type specific ARF functions. In future studies, it will be particularly important to pin-point the role of ARFs in polarized cell types and organ-like model systems due to their more complex membrane organization and to the presence of multiple segregating secretory routes.

ARF1 and ARF3 define TGN-derived tubular trafficking intermediates positive for clathrin (Figure7). ARF3 differs from ARF1 in only seven amino acids at the very C-and N-termini and the specific localization of ARF3 to the TGN has been mapped to two single C terminal residues (A174 and K180) conserved across species (Manolea et al., 2010). Previous reports highlighted differential effects on membrane recruitment of ARF1 and ARF3 when ARF GEFs of the BIG family are depleted (Manolea et al., 2010). Additionally, differential effects on trafficking when ARF1 and ARF3 are knocked down in combination with other ARFs have been observed (Volpicelli-Daley et al., 2005), strongly suggesting non-overlapping roles in TGN export for type I ARFs. Here, using live-cell STED microscopy, we show that gene edited ARF1 and ARF3 localize to segregating nano-domains on the TGN and downstream TGN-derived trafficking intermediates (Figure 8). This differential localization strengthens the idea that ARF1 and ARF3 may have differential sorting functions. The presence of ARF1 and ARF3 on the same TGN-derived tubules could lead to segregation of cargoes for sorting to downstream endosomal compartments. Interestingly, gene edited ARF3 with a GC C-terminal linker fails to correctly localize to the Golgi and post-Golgi carriers and shows very high cytoplasmic background (Supplementary figure 1). The addition of a long linker in the form of a LAP tag could possibly free the very C terminus of ARF3 for interaction with important uncharacterized membrane recruiting factors.

Lastly it is interesting to note that the higher numbers of ARF isoforms in mammals compared to yeast seems to be correlated with the more complex membrane organization of mammalian cells. Yeast does not possess ERGICs and have less complex post-Golgi secretory routes. Defining the function of this important classes of transport regulators will have broader implications on our understanding of the organization of intracellular membranes and cellular membrane homeostasis.

## Materials and methods

### Cell culture

HeLa cells ATCC CCL-2 (ECACC General Collection) and HAP1 cells (Essletzbichler et al., 2014) were grown at 37 °C with 5% CO_2_ in Dulbecco’s Modified Eagle Medium (DMEM, Gibco) supplemented with 10% fetal bovine serum (FBS, Corning), 100 U/L penicillin and 0.1 g/L streptomycin (FisherScientific).

Transfection of plasmid DNA in HeLa cells was carried out using FuGENE (Promega). For the DNA transfection, 2 μg of plasmid DNA and 6 µL of FuGENE per well were used. Transfection of plasmid DNA in HAP1 cells was carried out using a NEPA21 electroporation system (Nepa Gene). Approximately 3 million cells were washed twice with Opti-MEM (Gibco) and then resuspended in 90 µL Opti-MEM with 10 µg of DNA in an electroporation cuvette with a 2 mm gap. HAP1 cells were electroporated with a poring pulse of 200 V, 5 ms pulse length, 50 ms pulse interval, 2 pulses, with decay rate of 10% and + polarity; consecutively with a transfer pulse of 20 V, 50 ms pulse length, 50 ms pulse interval, 5 pulses, with a decay rate of 40% and ± polarity.

Nocodazole (Sigma), G418 (Gibco) and puromycin (Gibco) were used at the following concentrations: nocodazole 2.5 μg/ml; G418 1 mg/ml (HeLa cells) or 3 mg/ml (HAP1 cells); and puromycin 1 μg/ml (HeLa cells).

### Preparation of cells for live-cell imaging

For all of the live-cell imaging experiments, cells were seeded on a glass-bottom dish (3.5 cm diameter, No. 1.5 glass; Cellvis) coated with fibronectin (Sigma). Labeling with Halo and SNAP substrates for imaging was carried out for 1 h using 1 μM stocks. After the staining, cells were washed 3 times with growth medium to get rid of the excess of dye and left for 1 h in an incubator at 37 °C and 5% CO_2_. To break down the Golgi ribbon into ministacks, cells were first labeled and subsequently treated with nocodazole for 3 h. All live-cell imaging were carried out in FluoBrite DMEM (Gibco) supplemented with 10% FBS, 20 mM HEPES (Gibco) and 1x GlutaMAX (Gibco). All live-cell experiments were carried out at 37°C. See Supplementary Table 1 for the list of dyes used.

### Molecular biology and CRISPR-Cas9 gene editing

See the supplementary methods.

### Immunofluorescence

Immunofluorescence was carried out on cells seeded on 1.5# glass 12 mm coverslips previously coated with fibronectin. All steps were carried out at room temperature (RT). First, samples were rinsed three times with PBS and then fixed using 4 % PFA (16 % stock, Electron Microscopy Sciences) for 10 min. After fixation, samples were rinsed three times with PBS and then permeabilized using permeabilization buffer (0.3 % NP40, 0.05 % Triton-X 100 and 0.1 % BSA (IgG free) in PBS). Cells were rinsed three times with wash buffer (0.05 % NP40, 0.05 % Triton-X 100 and 0.2 % BSA in PBS) and blocked for 45 min in block buffer (0.05 % NP40, 0.05 % Triton-X 100 and 5 % goat serum (Jackson ImmunoResearch) in PBS). Subsequently, cells were incubated with primary antibodies (1:1000) for 1 h in block buffer and then washed three times for 5 min with wash buffer. Secondary antibody (1:2000) incubation was carried out for 30 min in block buffer. Again, samples were washed three times for 5 min with wash buffer and then rinsed three times with PBS. Samples were then dipped in ddH_2_O and mounted with pro-long Gold (Life Technologies). See Supplementary Table 2 for the list of antibodies used.

### STED imaging

STED imaging was performed on a commercial expert line Abberior STED microscope equipped with 485 nm, 561 nm, and 645 nm excitation lasers. The 775 nm depletion laser was used to deplete dyes in two color STED experiments. Two-color images were recorded sequentially line by line. Detection was carried out with avalanche photodiodes (APDs) with detection windows set to 571-630 nm and 650-756 nm. When required, excitation power was kept constant between samples to allow the quantification of signal differences. The pixel size was set to 60 nm for confocal imaging, 30 nm for live-cell STED imaging, and 20 nm for STED imaging in fixed cells.

### Image processing and statistical analysis

Raw STED images were deconvolved to reduce noise using Richardson–Lucy deconvolution from the python microscopy PYME package (https://python-microscopy.org/), background subtracted and smoothed using a Gaussian filter with 1-pixel SD using ImageJ (Abramoff, 2004) as indicated in the figures.

Mean fluorescence intensities at the Golgi (Figure 1j) were measured using ImageJ. To select the area for quantification of the signal intensity at the Golgi, a mask of the Golgi was determined by a thresholding method from ImageJ (MaxEntropy).

Line profile data shown in Figure 2,3,4,6,8 and Supplementary figure 4,5,8 was obtained from gaussian blurred images using ImageJ by drawing a line perpendicular to the direction of the cisternae. Line profile data from ImageJ was plotted using Graphpad Prism.

The quantification of the number of ARF1-and ARF4-positive distal ERGICs (Figure 4d) was done using raw image data in ImageJ and plotted using Graphpad Prism. Representative images are presented in Supplementary figure 7. The percentage of distal ERGIC positive for either ARF1 or ARF4 was calculated by counting all ARF1/ARF4 positive distal ERGICs divided by the total amount of ERGICs in ARF1^EN^-and ARF4^EN^-Halo KIs cells expressing SNAP-ERGIC53. The percentage of ARF1-and ARF4-positive distal structures was calculated by counting all ARF1 and ARF4 positive structures divided by the total amount of distal ARF4 structures in ARF1^EN^-SNAP and ARF4^EN^-Halo double KI cells.

To quantify the distance from the edge of the ARF-labelled cisternae to the GM130-labelled *cis*-Golgi cisternae (Figure 3j) and to the Golgin97-labelled TGN cisternae (Figure 6f) in nocodazole-treated fixed ARFs^EN^-2xALFA KI cells, a box perpendicular to the direction of the cisternae was drawn to obtain a line profile using ImageJ. The box profile data was further analyzed with a custom Python code (available on https://github.com/AG-Bottanelli/distance-to-edge.git) that defined the edge of the cisternae. The edge of each cisterna (ARFs, GM130, Golgin97 and ERGIC53) was calculated as the coordinate with a maximum mean intensity gradient. After the first computation attempt, we plotted the line profiles alongside with their gradients and discarded those in which the edge was detected in a second peak of signal for M shaped profiles. The data obtained from the Python script was plotted using Graphpad Prism.

To calculate the distance from the edge of the ARF-labelled cisternae to the center of COPI-positive (Figure 5d) or clathrin-positive (Figure 7f) clusters in live nocodazole-treated ARFs^EN^-Halo+BetaCOP^EN^-SNAP/SNAP-CLCa^EN^ double KI cells, line profiles were obtained by drawing a box between the ARF cisternae and the vesicle using ImageJ. Again, the line profile data was further analyzed with a Python script (available on https://github.com/AG-Bottanelli/distance-to-edge.git) that defined the center of a COPI/clathrin cluster and the edge of the cisternae as described previously. The center of COPI/clathrin clusters was determined as the position of the maximum mean signal intensity value. After the first computation attempt, we plotted the line profiles alongside with their gradients and discarded those in which (1) “vesicle” signal’s FWHM was higher than 200 nm or (2) edge was detected in a second peak of signal for M shaped profiles. The data obtained from the Python script was plotted using Graphpad Prism.

The quantification of the mean COPI and clathrin signal at the Golgi in HAP1 cells (Supplementary Figure 2c-d) was measured using raw data in ImageJ. To select the area for quantification of the signal intensity at the Golgi, a mask of the Golgi was determined by using the thresholding function (Moments) from ImageJ. HAP1 WT and HAPI ARFs^EN^-Halo KI IF samples were prepared and imaged on the same day using the same parameters on the STED microscope to ensure comparability between samples. Signal intensities for quantification come from three independent experiments. The signal intensities from the COPI/clathrin signal in the ARFs^EN^-Halo KI cells were normalized to the mean signal intensity from the WT cells.

### SDS-PAGE and Western Blot

Protein lysates of HAP1 and HeLa cells were separated by sodium dodecyl sulfate polyacrylamide gel electrophoresis (SDS-PAGE) using 4-12% gradient precast polyacrylamide gels (Life Technologies). The proteins separated by SDS-PAGE were transferred onto a nitrocellulose membrane using a Mini-Protean® Tetra Cell device. Halo tag fusions were detected using a primary anti-Halo antibody at a dilution of 1:1000 in PBS supplemented with 1 % BSA and 0.02 % sodium azide. For the detection of primary antibodies, anti-mouse horseradish peroxidase-coupled (1:5000) secondary antibodies were used. For final detection, Pierce ECL substrate was used, and bioluminescent signal was detected.

## Acknowledgments

We would like to thank the whole lab for the many discussions and for reading this manuscript. A particular thank you goes to Blanche Schwappach and Andreas Ernst for critically reading the manuscript and for the many engaging discussions on the topic. We thank Jez Carlton for suggesting the use of the LAP tag. We thank Luke Lavis, Jonathan Grimm and the team of chemists at Janelia that provide us with the latest and brightest STEDable dyes for our experiments. This project was supported by Deutsche Forschungsgemeinschaft (DFG, German Research Foundation) grants SFB958 (Project A25 (C.R. and A.S.)), SFB/TRR186 (Project A20 (L.W-D. and P.A.)) and a major research instrumentation grant for the acquisition of the STED microscope.

