## Supplementary material for "Gene editing and super-resolution microscopy reveal multiple distinct roles for ARF GTPases in cellular membrane organization"

### Supplementary figures

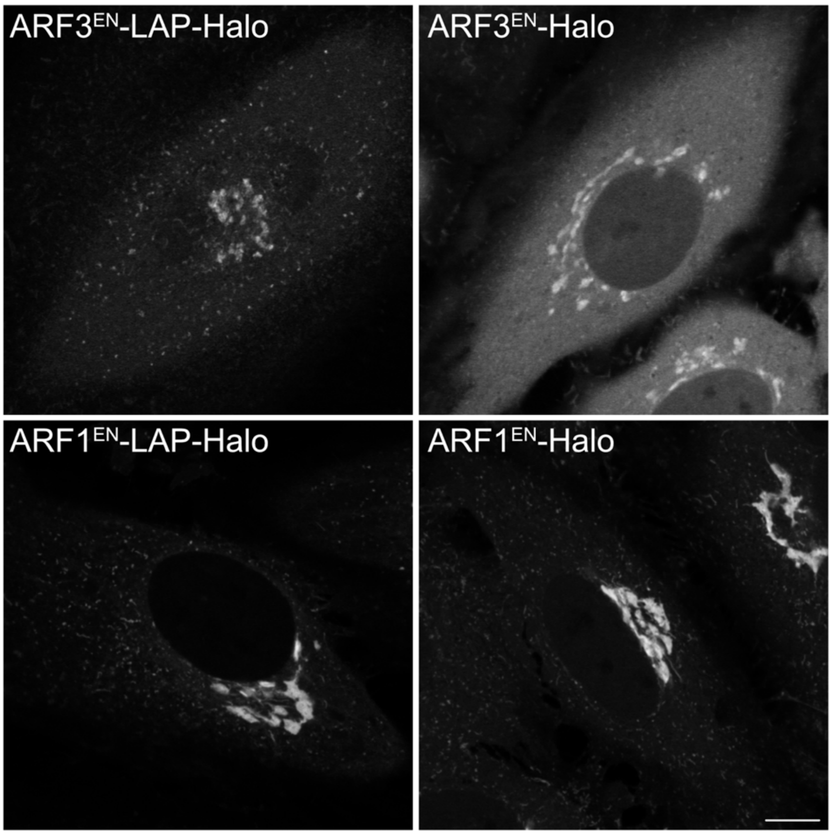


**Supplementary figure 1. Addition of a LAP tag greatly improves the localization of endogenously tagged ARF3.** Endogenously tagged ARF GTPases either with a GS linker (ARF1$^{EN}$-Halo and ARF3$^{EN}$-Halo) or with a LAP tag linker where GFP was switched with Halo tag (ARF1$^{EN}$-LAP-Halo and ARF3$^{EN}$-LAP-Halo) labelled with JFX650-CA. While addition of a LAP linker improves the cytoplasmic localization of ARF3, no differences were observed in the localization of ARF1 with or without a LAP linker. All images were smoothed with a gaussian filter as described in the methods. CA=chloroalkane. Scale bars are 10 µm.

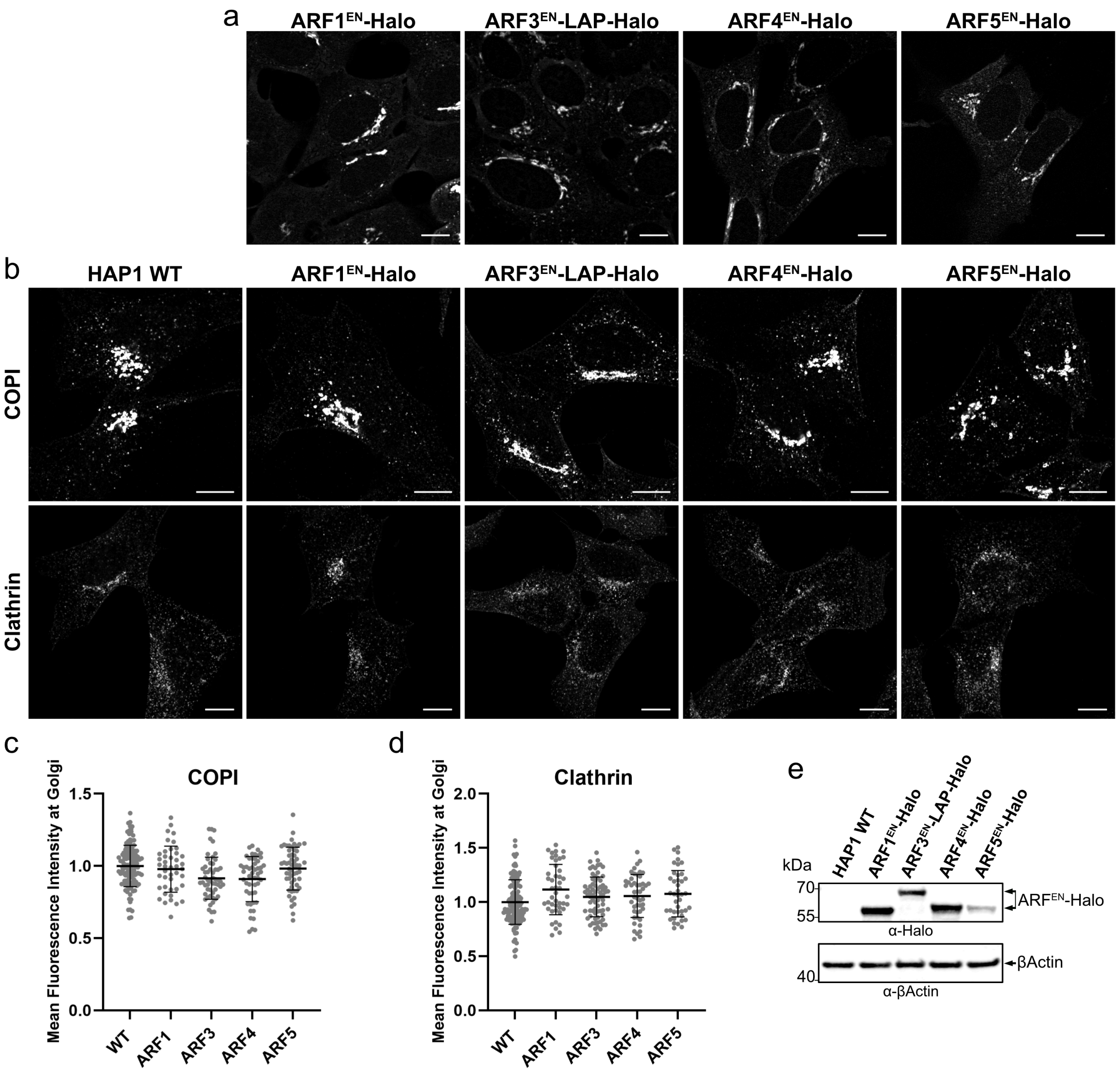


**Supplementary figure 2. HAP1 cells edited at each ARF locus do not present any morphological defects or defects in coats recruitment.** ARFs were tagged at their endogenous locus with the self-labelling enzyme Halo in a HAP1 haploid cell line. Live cells were stained with the Halo substrate JFX650-CA (a). HAP1 wild type (WT) and ARF$^{EN}$-Halo KI cells were immunostained with anti-COPI and anti-clathrin and secondary antibodies conjugated to Alexa488 (b). Scatter dot plots with mean and standard deviation represent the quantification of the mean fluorescence intensity of COPI (c) and clathrin (d) at the Golgi measured in HAP1 WT and ARF$^{EN}$-Halo KI cells as described in the methods. Immunoblot to detect ARF-Halo fusion proteins in HAP1 KI lysate with anti-Halo antibodies and anti-βActin primary antibody as loading control (e). All images were smoothed with a gaussian filter as described in the methods. Scale bars are 10 µm.

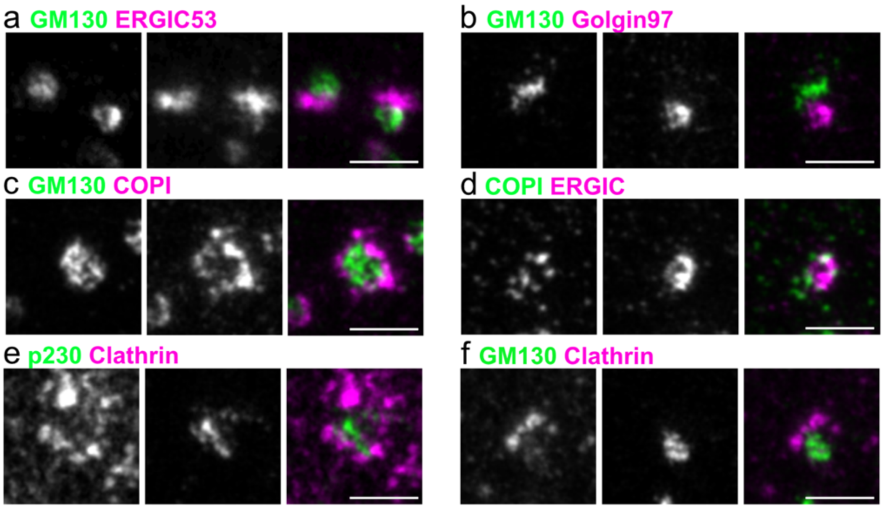


**Supplementary figure 3. Mapping of Golgi markers with STED microscopy in nocodazole treated cells.** HeLa cells were treated with nocodazole (2.5 µg/mL) for 3h, fixed and immunostained as indicated in the figure. Secondary antibodies labelled either with ATTO647N or AlexaFluor594 were used to be able to perform dual-color STED experiments. All images were smoothed with a gaussian filter as described in the methods. Scale bars are 1 µm.

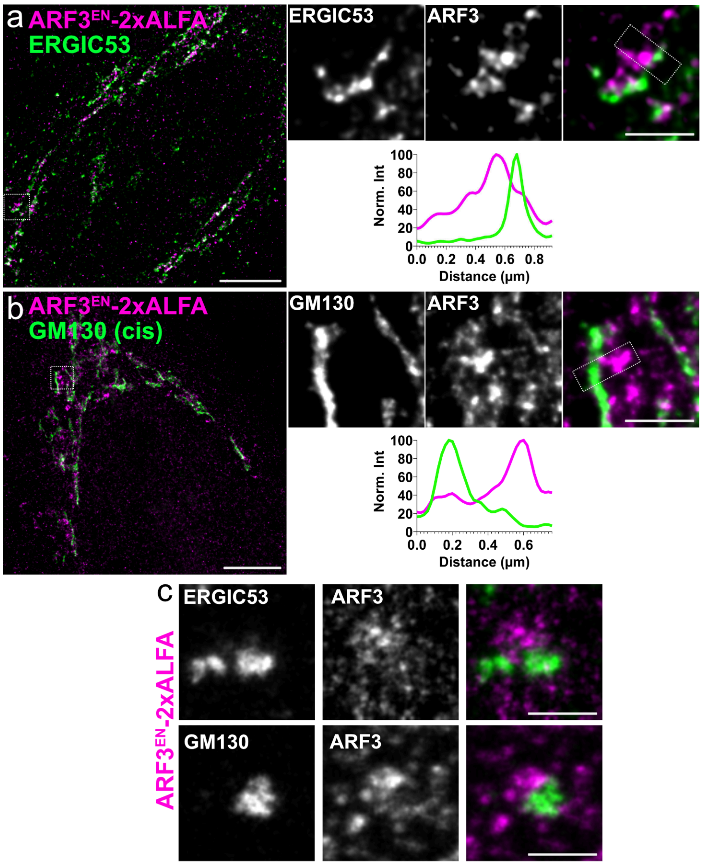


**Supplementary figure 4. STED microscopy reveals ARF3 clusters segregated from ERGIC and *cis*-Golgi cisternae.** ARF3$^{EN}$-2xALFA (magenta) KI HeLa cells were fixed and immunostained with anti-ALFA tag and either anti-ERGIC53 (a, green) or anti-GM130 (b, green). Secondary antibodies labelled with either ATTO647N or AlexaFluor594 were used to be able to perform dual-color STED experiments (b). ARF3$^{EN}$-2xALFA KI HeLa cells were treated with nocodazole (2.5 µg/mL) for 3h and immunostained with the same primary antibodies. Secondary antibodies labelled with either ATTO647N or AlexaFluor594 (GM130)/AlexaFluor568 (ERGIC53) were used (c). All images were deconvolved and smoothed as described in the methods. Line profiles in each panel correspond to the dotted boxes in the cropped images. Scale bars are 5 µm and 1 µm in the cropped images.

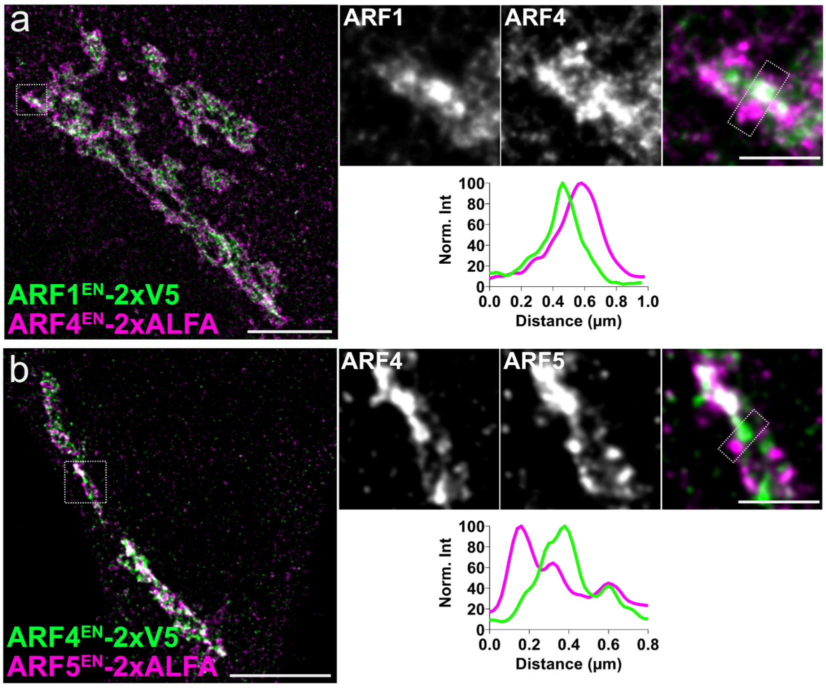


**Supplementary figure 5. STED microscopy of double edited ARF1-ARF4 and ARF4-ARF5 cells suggests distinct functions.** ARF1$^{EN}$-2xV5 (a, green)/ARF4$^{EN}$-2xALFA (a, magenta) and ARF4$^{EN}$-2xV5 (b, green)/ARF5$^{EN}$-2xALFA (b, magenta) double KI HeLa cells were fixed and immunostained with anti-ALFA or anti-V5 primary antibodies and secondary antibodies labelled with either ATTO647N or AlexaFluor594 to be able to perform dual-color STED experiments. Line profiles in each panel correspond to the dotted boxes in the cropped images. Scale bars are 5 µm and 1 µm in the cropped images.

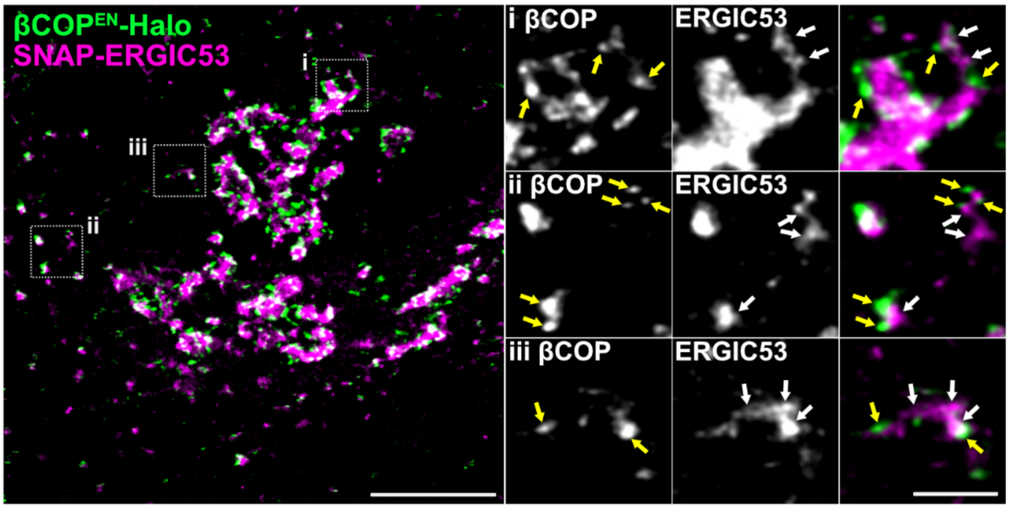


**Supplementary figure 6. Live-cell STED highlights tubular vesicular ERGIC elements decorated with COPI clusters.** βCOP$^{EN}$-Halo (green) KI HeLa cells were transfected with a plasmid encoding for SNAP-ERGIC53. Live cells were stained with the Halo substrate JF571-CA together with the SNAP substrate JFX650-BG. Yellow arrows highlight COPI-positive clusters associated with tubular-vesicular ERGIC elements (white arrows). CA=chloroalkane, BG= benzylguanine. Scale bars are 5 µm and 1 µm in the cropped images.

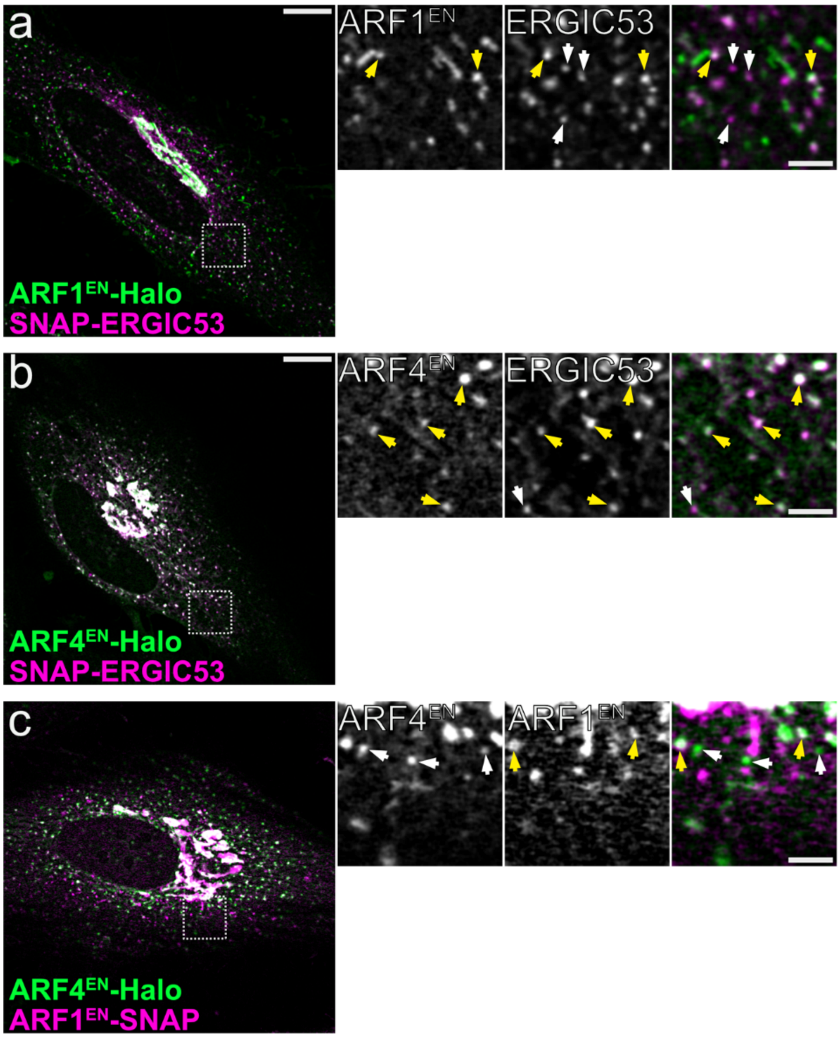


**Supplementary figure 7. ARF1 and ARF4 define different sub-populations of distal ERGICs.** ARF1$^{EN}$-Halo (a) and ARF4$^{EN}$-Halo (b) KI cells were transfected with a plasmid encoding for SNAP-ERGIC53 and labelled with the Halo substrate JF552-CA and the SNAP substrate JFX650-BG. White arrows highlight ARF1- or ARF4-positive only structures while yellow arrows highlight structures in which ARFs are co-localizing with ERGIC53. Double KI ARF4$^{EN}$-Halo/ARF1$^{EN}$-SNAP (c) were labelled with the Halo substrate JF552-CA and the SNAP substrate JFX650-BG. White arrows highlight ARF4 positive structures while yellow arrows highlight structures were ARF1 and ARF4 co-localize. CA=chloroalkane, BG= benzylguanine. Scale bars are 10 μm and 2 μm in the cropped images.

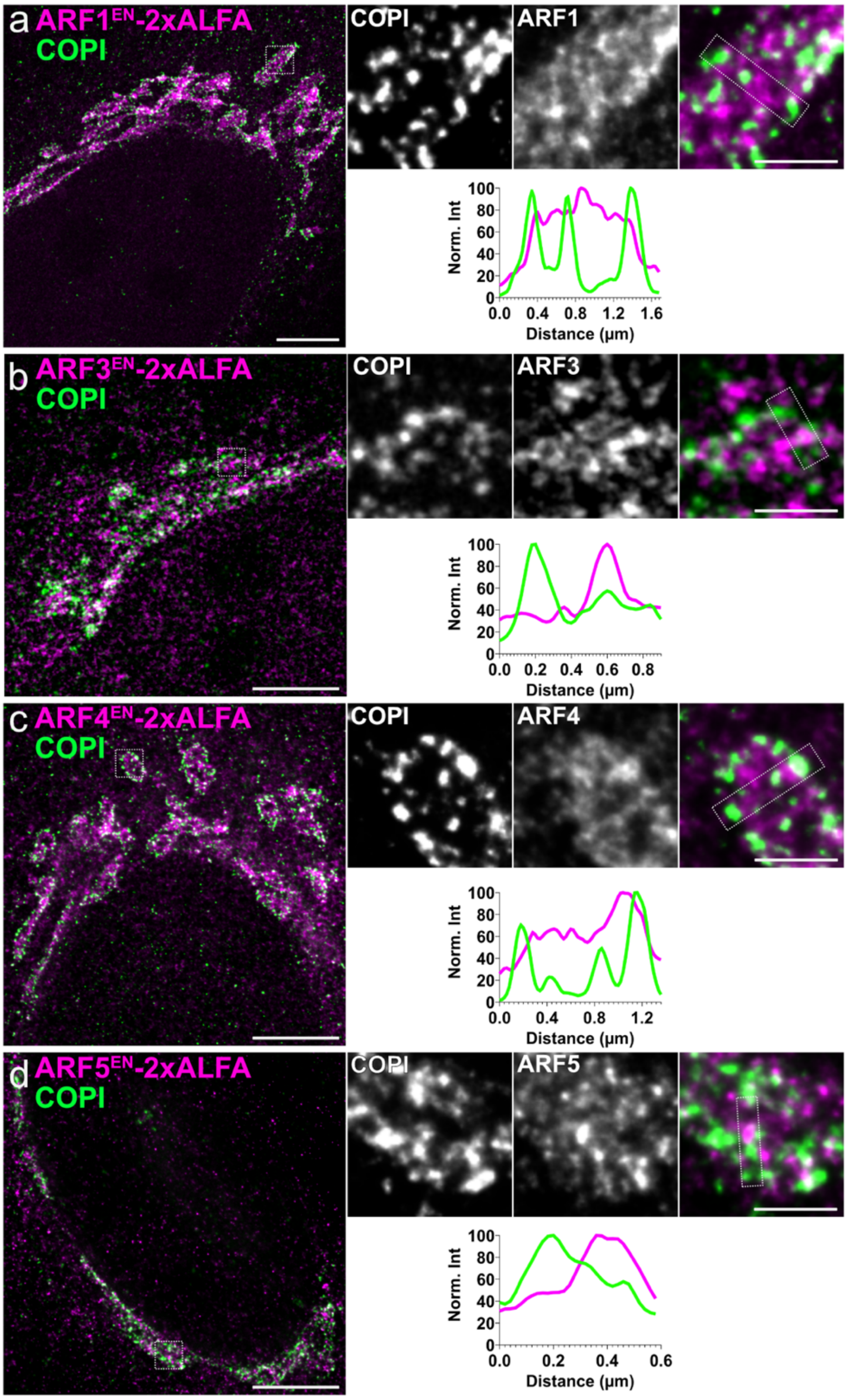


**Supplementary figure 8. COPI-positive clusters are observed near all ARFs.** Fixed $ARF^{EN}$-2xALFA KIs cell lines were immunostained with anti-ALFA (magenta) and anti-β'COP (CM1, green) and secondary antibodies labelled with either ATTO647N or AlexaFluor594 to be able to perform dual-color STED experiments. Line profiles in each panel correspond to the dotted boxes in the cropped images. Scale bars are 5 µm and 1 µm in the cropped images.

### Supplementary methods

#### Recombinant Plasmids for overexpression

To generate the SNAP-ERGIC53 plasmid, a pC4 vector was linearized with EcoRI and BamHI and SNAP and ERGIC53 fragments were inserted as EcoRI-HindIII and HindIII-BamHI fragments, respectively. SNAP was amplified with SNAP-SPs-EcoRI-sense 5’-TACGTGAATTCATGAGGTCTTTGCTAATCTTGGTGCTTTGCTTCCTGCCCCTGGCTGCTCTGGGGAAAGACAAAGACTGCGAAATGAAGCGC-3’ and SNAP-HindIII-anti-sense 5’-ACGATCAAGCTTTCCTGAGCCACCCAGCCCAGGCTTGCCCAGTCTGTGGCCC-3’. The SNAP sense oligo also contains a short sequence encoding for chicken lysozyme signal peptide (SP). The sequence encoding for ERGIC53 was amplified from YFP-ERGIC53 (Hamlin et al., 2014) with ERGIC53-HindIII-sense primer 5’-GATCGTAAGCTTTACTCAGGAGGGGGTGGCGACGGCATGGGAGGAGATGCTGCGGCT-3’ and ERGIC53-BamHI-anti-sense 5’-CAGTACGGATCCTCAGAAGAATTTTTTGGCAGCTGCTTCTTGCTGAGTCCTGTACA-3’.

#### CRISPR-Cas9 Gene Editing

The ARF1 genomic locus (gene ID 375) was targeted as previously described (Bottanelli et al., 2017). The ARF3 genomic locus (gene ID 377) was targeted with the guide RNA: 5’- GGTGCTTTGTTAGGGCTGTCTGG-3’ (PAM sequence is underlined) located after the stop codon in the ARF3 coding sequence. The ARF4 genomic locus (gene ID 378) was targeted with the guide RNA: 5’-TGAAATTGGATATCTAACCAAGG-3’ located after the stop codon in the ARF4 coding sequence. The ARF5 genomic locus (gene ID 381) was targeted with the guide RNA: 5’-TCAAAGCGCTAACCAGCCAGGGG-3’ located after the stop codon in the ARF5 coding sequence. The COPB1 gene (gene ID 1315) was targeted with the guide RNA: 5’-AAATTAACTGTAAAGCTTCAAGG-3’ located after the stop codon in the COPB1 coding sequence. The CLTA gene (gene ID 1211) was targeted using the guide RNA: 5’-ATGGCTGAGCTGGATCCGTTCGG-3’ located after the start codon in the CLTA coding sequence. All guide RNAs were designed using Benchling (https://www.benchling.com) and no guide RNAs have significant off-target matches when checked with the CRISPR design tool. The guide RNAs were cloned into the pSpCas9(BB)-2A-Puro (pX459) V2.0 (Addgene plasmid #62988) (Ran et al., 2013) by annealing oligoes and ligating them into the vector previously linearized with BpiI (Thermo Fisher). The guide RNA to target the CLTA locus was cloned into the SpCas9 pX330 plasmid (Addgene plasmid #42230) (Cong et al., 2013) vector using the same strategy described above.

The homology repair (HR) plasmid for ARF1-Halo/SNAP was previously described in Bottanelli et. al., 2017. The HR plasmid to generate gene edited ARF3-Halo, ARF4-Halo/SNAP and ARF5-Halo were constructed starting from pTWIST Amp High Copy plasmids (Origin: pMB1). The DNA sequence of the left and right homology arms (ca. 1 Kb each) from ARF3 and ARF5 were synthesized by Twist Bioscience with NheI and BamHI sites to insert the desired tag and a glycin/serin rich linker (GSGSGSGSGS). In the case of ARF4, only the the right homology arm sequence was synthesized by Twist Bioscience with EcoRI, NheI and BamHI sites for further downstream manipulations. The ARF4 right homology arm was amplified from genomic DNA using the following oligoes: ARF4-Left-Homolgy-Arm-sense 5'-GTCATGAATTCCTGCGTTAGCCAGGAAGGTCTCGATC-3' and ARF4-Left-Homology-Arm-anti-sense 5'-TCATACGCTAGCTGAGCCGGAACCAGAGCCTGACCCTGATCCACGTTTTGAAAGCTCATTTGACAGCCAGTCAAGTC-3' and cloned as an EcoRI-NheI fragment. Halo/SNAP tags were introduced into pTWIST ARF3/ARF4/ARF5 homology repair plasmids as NheI-BamHI fragments. The Halo tag sequence was amplified out of the pHalo-N1 ARF1-Halo homology repair plasmid (Bottanelli et al., 2017) using the following primers: NheI-Halo-sense 5'-CTGATGCTAGCATGGCAGAAATCGGTACTGGCTTTC-3' and Halo-BamHI-anti-sense 5'-TACGAGGATCCTCAAGCGTAATCTGGAACATCGTATGGGTAAGCGTAATCTGGAACATCGTATGGGTAGCCGGAAATCTCGAGCGTCGACAGCCAGC-3'. The SNAP tag sequence was amplified out of the pSNAPf plasmid (New England Biolabs) using the following primers: NheI-SNAP-sense 5'-CTGATGCTAGCGACAAAGACTGCGAAATGAAGCGCA-3' and SNAP-BamHI-anti-sense 5'-TACGAGGATCCTCACTTGTCGTCATCGTCTTTGTAGTCCTTGTCGTCATCGTCTTTGTAGTCACCCAGCCCAGGCTTGCCCAGTCTGTGGCC-3'. The PAM site in all HR plasmids was mutagenized to prevent cutting by the Cas9.

To facilitate selection of positive recombinants with the drug G418/Geneticin (Gibco), the $Neo^r$/$Kan^r$ resistance cassette flanked by LoxP sites was introduced into the ARF3/ARF4/ARF5-Halo HR plasmids as a BamHI-BamHI fragment. This allows selection of the positively edited cells with G418 and subsequent excision of the resistance cassette to restore the endogenous locus (Bottanelli et al., 2017; Schmidt et al., 2016). The LoxP-$G418^r$-LoxP fragment was amplified with the following oligoes LoxP-$G418^r$-sense 5'-ATGTCGGATCCATAACTTCGTATAGCATACATTATACGAAGTTATCCTGAGGCGGAAAGAACCAGCTGTGGAATGTGTGTCAGTTAG-'3 and $G418^r$-LoxP-anti-sense 5'-TGCACGGATCCATAACTTCGTATAATGTATGCTATACGAAGTTATTTTATTCTGTCTTTTTATTGCCGTCATAGCGCGGGTT-'3 from a pEGFP-N1 vector (Clontech).

The HR plasmid to generate gene edited ARF3/ARF4/ARF5-2xALFA were generated by linearizing the respective ARF-Halo HR plasmid with NheI and BamHI sites. To generate the ARF1-2xALFA HR plasmid, a new HR plasmid was synthesized (Twist Bioscience) to introduce NheI and BamHI sites and a glycin/serin rich linker in the same way as previously designed for ARF3 and ARF5. The sequence encoding for 2xALFA tag was synthesized as a double strand DNA fragment (Integrated DNA technologies) and assembled into the linearized NheI-BamHI ARFs HR plasmids via Gibson assembly. In case of ARF1 and ARF3, a new EcoRI site was introduced between the 2xALFA tag and the BamHI site to facilitate further manipulations and allow the excision of the tag or the resistance cassette independently. In case of ARF4 and ARF5, a new HindIII site was introduced between the 2xALFA tag and the BamHI site. The following double strand DNA fragments were used:

ARF1-NheI-2xALFA-EcoRI-BamHI: 5'-TGGCTGTCCAATCAGCTCCGGAACCAGAAGGGATCAGGGTCAGGCTCTGGTTC

CGGCTCAGCTAGCAGCAGACTGGAGGAGGAGCTGAGAAGAAGACTGACCGAG GGCAGCCGTCTGGAAGAAGAACTGCGTCGTCGTCTGACCGAATGAGAATTCAT CGTAGCGGATCCACGCGACCCCCCTCCCTCTCACTCCTCTTG-3’

ARF3-NheI-2xALFA-EcoRI-BamHI: 5’-TGGCTGGCCAATCAGCTCAAAAACAAGAAGGGATCAGGGTCAGGCTCTGGTTC CGGCTCAGCTAGCAGCAGACTGGAGGAGGAGCTGAGAAGAAGACTGACCGAG GGCAGCCGTCTGGAAGAAGAACTGCGTCGTCGTCTGACCGAATGAGAATTCAT CGTAGCGGATCCAAGCGAGACAGCGCTAACAAAGCACCCCAC-3’

ARF4-NheI-2xALFA-HindIII-BamHI: 5’-GACTGGCTGTCAAATGAGCTTTCAAAACGTGGATCAGGGTCAGGCTCTGGTTCC GGCTCAGCTAGCAGCAGACTGGAGGAGGAGCTGAGAAGAAGACTGACCGAGG GCAGCCGTCTGGAAGAAGAACTGCGTCGTCGTCTGACCGAATGAAAGCTTATC GTAGCGGATCCATGAAATTCGATATCTAACCAACGACATGT-3’

ARF5-NheI-2xALFA-HindIII-BamHI: 5’-GACTGGCTGTCCCACGAGCTGTCAAAGCGCGGATCAGGGTCAGGCTCTGGTTC CGGCTCAGCTAGCAGCAGACTGGAGGAGGAGCTGAGAAGAAGACTGACCGAG GGCAGCCGTCTGGAAGAAGAACTGCGTCGTCGTCTGACCGAATGAAAGCTTAT CGTAGCGGATCCCCAGCGAGGCGCAGGCCCCTGATGCCCGGA-3’

Again, to facilitate selection of positive recombinants with G418, the Neo$^r$/Kan$^r$ resistance cassette flanked by LoxP sites was introduced into the ARF1/ARF3/ARF4/ARF5-2xALFA HR plasmids as an EcoRI-BamHI (ARF1 and ARF3) or an HindIII-BamHI (ARF4 and ARF5). The EcoRI/HindIII-LoxP-G418$^r$-LoxP-BamHI fragments were amplified from a pEGFP-N1 vector with the following oligoes: EcoRI/HindIII-LoxP-G418$^r$-sense 5’-ATGTC(GAATTC/AAGCTT)ATAACTTCGTATAGCATACATTATACGAAGTTATCCT GAGGCGGAAAGAACCAGCTGTGGAATGTGTGTCAGTTAG-3’ and G418$^r$-LoxP-BamHI-anti-sense 5’-TGCACGGATCCATAACTTCGTATAATGTATGCTATACGAAGTTATTTTATTCTGTC TTTTTATTGCCGTCATAGCGCGGGTT-3’.

The ARF1/ARF4-2xV5 HR plasmids were generated using the same strategy as for the ARFs-2xALFA HR plasmids using Gibson assembly. The following double strand DNA fragments were inserted into the NheI-BamHI linearized ARF HR plasmids:

ARF1-NheI-2xV5-EcoRI-BamHI: 5’-TGGCTGTCCAATCAGCTCCGGAACCAGAAGGGATCAGGGTCAGGCTCTGGTTC CGGCTCAGCTAGCGGCAAGCCCATCCCCAACCCCCTGCTGGGCCTGGACAGC ACCGGCGGTAAACCAATTCCAAATCCATTGTTGGGTTTGGATTCTACTTGAGAAT TCATCGTAGCGGATCCACGCGACCCCCCTCCCTCTCACTCCTCTTG-3’

ARF4-NheI-2xV5-HindIII-BamHI: 5’-GACTGGCTGTCAAATGAGCTTTCAAAACGTGGATCAGGGTCAGGCTCTGGTTCC GGCTCAGCTAGCGGCAAGCCCATCCCCAACCCCCTGCTGGGCCTGGACAGCA CCGGCGGTAAACCAATTCCAAATCCATTGTTGGGTTTGGATTCTACTTGAAAGCT TATCGTAGCGGATCCATGAAATTCGATATCTAACCAACGACATGT-3’

To select for positively gene edited cells, the sequence of the Puromycin resistance cassette (Puromycin$^{r}$) flanked by LoxP sites was introduced into the ARF1/ARF4-2xV5 HR plasmids linearized with EcoRI/HindIII-BamHI. The EcoRI/HindIII-LoxP-Puromycin$^{r}$-LoxP-BamHI fragments were amplified from a HindIII-mutagenized pPUR vector using the following primers: EcoRI/HindIII-LoxP-Puro$^{r}$-sense 5'-ATGTC(<u>GAATTC/AAGCTT</u>)ATAACTTCGTATAGCATACATTATACGAAGTTATCTG
TGGAATGTGTGTCAGTTAGGGTGTGGAAAGTCCCCAGGCTCCC-3' and Puro$^{r}$-LoxP-BamHI-anti-sense 5'-
TGCACGGATCCATAACTTCGTATAATGTATGCTATACGAAGTTATCAGACATGAT
AAGATACATTGATGAGTTTGGACAAACCACAAC-3'. The HindIII site in the Puromycin$^{r}$ cassette in the pPUR vector (Clonetech) was mutagenized with the following oligoes: Puro$^{r}$-HindIII-G342C-sense 5'-CTAGGCTTTTGCAAAAACCTTGCATGCCTGCAGGTC-3' and Puro$^{r}$-HindIII-G342C-anti-sense: 5'-GACCTGCAGGCATGCAAGGTTTTTGCAAAAGCCTAG-3'.

ARF3-LAP-Halo-2xALFA-LoxP-G418$^{r}$-LoxP HR and ARF3-LAP-SNAP-2xV5-LoxP-Puromycin$^{r}$-LoxP HR plasmids were generated by linearizing the ARF3-2xALFA-LoxP-G418$^{r}$-LoxP HR and ARF3-2xV5-LoxP-Puromycin$^{r}$-LoxP HR with NheI. The LAP-Halo tag and LAP-SNAP tag were synthesized as double stranded DNA fragments and assembled into the respective NheI linearized ARF3-2xALFA-LoxP-G418$^{r}$-LoxP HR and ARF3-2xV5-LoxP-Puromycin$^{r}$-LoxP HR plasmids via Gibson assembly.

The following double strand DNA fragments were inserted into the NheI linearized ARF3 HR plasmids:

LHA-GS-NheI-LAP-Halo-2xALFA: 5'-
TGGCTGGCCAATCAGCTCAAAAACAAGAAGGGATCAGGGTCAGGCTCTGGTTC
CGGCTCAGCTAGCATGTACCCAACTTTCTTGTACAAAGTGGTACCGGTCGCCAC
CGCAAGCGAGAATCTTTATTTTCAGGGCGCCGCCAAATTCAAAGAAACCGCTGC
TGCTAAATTCGAACGCCAGCACATGGACAGCGGAGGTGGAGGTTCAAGTGGTC
CCTCAGGTTCGTCGAGCCTGGAAGTTCTGTTCCAGGGGCCCCTGTCGTCTAGC
GGTCCCTCGGGTTCTATGGCAGAAATCGGTACTGGCTTTCCATTCGACCCCCAT
TATGTGGAAGTCCTGGGCGAGCGCATGCACTACGTCGATGTTGGTCCGCGCGA
TGGCACCCCTGTGCTGTTCCTGCACGGTAACCCGACCTCCTCCTACGTGTGGC
GCAACATCATCCCGCATGTTGCACCGACCCATCGCTGCATTGCTCCAGACCTGA
TCGGTATGGGCAAATCCGACAAACCAGACCTGGGTTATTTCTTCGACGACCACG
TCCGCTTCATGGATGCCTTCATCGAAGCCCTGGGTCTGGAAGAGGTCGTCCTG
GTCATTCACGACTGGGGCTCCGCTCTGGGTTTCCACTGGGCCAAGCGCAATCC
AGAGCGCGTCAAAGGTATTGCATTTATGGAGTTCATCCGCCCTATCCCGACCTG
GGACGAATGGCCAGAATTTGCCCGCGAGACCTTCCAGGCCTTCCGCACCACCG
ACGTCGGCCGCAAGCTGATCATCGATCAGAACGTTTTTATCGAGGGTACGCTGC
CGATGGGTGTCGTCCGCCCGCTGACTGAAGTCGAGATGGACCATTACCGCGAG
CCGTTCCTGAATCCTGTTGACCGCGAGCCACTGTGGCGCTTCCCAAACGAGCT
GCCAATCGCCGGTGAGCCAGCGAACATCGTCGCGCTGGTCGAAGAATACATGG
ACTGGCTGCACCAGTCCCCTGTCCCGAAGCTGCTGTTCTGGGGCACCCCAGGC
GTTCTGATCCCACCGGCCGAAGCCGCTCGCCTGGCCAAAAGCCTGCCTAACTG
CAAGGCTGTGGACATCGGCCCGGGTCTGAATCTGCTGCAAGAAGACAACCCGG
ACCTGATCGGCAGCGAGATCGCGCGCTGGCTGTCGACGCTCGAGATTTCCGGC
AGCAGACTGGAGGAGGAGCTGAGAAGAAGACTGACCGAGGGCAGCCGTCTG-3'

LHA-GS-NheI-LAP-SNAP-2xV5: 5’-
TGGCTGGCCAATCAGCTCAAAAACAAGAAGGGATCAGGGTCAGGCTCTGGTTC
CGGCTCAGCTAGCATGTACCCAACTTTCTTGTACAAAGTGGTACCGGTCGCCAC
CGCAAGCGAGAATCTTTATTTTCAGGGCGCCGCCAAATTCAAAGAAACCGCTGC
TGCTAAATTCGAACGCCAGCACATGGACAGCGGAGGTGGAGGTTCAAGTGGTC
CCTCAGGTTCGTCGAGCCTGGAAGTTCTGTTCCAGGGGCCCCTGTCGTCTAGC
GGTCCCTCGGGTTCTATGGACAAAGACTGCGAAATGAAGCGCACCACCCTGGA
TAGCCCTCTGGGCAAGCTGGAACTGTCTGGGTGCGAACAGGGCCTGCACCGTA
TCATCTTCCTGGGCAAAGGAACATCTGCCGCCGACGCCGTGGAAGTGCCTGCC
CCAGCCGCCGTGCTGGGCGGACCAGAGCCACTGATGCAGGCCACCGCCTGGC
TCAACGCCTACTTTCACCAGCCTGAGGCCATCGAGGAGTTCCCTGTGCCAGCC
CTGCACCACCCAGTGTTCCAGCAGGAGAGCTTTACCCGCCAGGTGCTGTGGAA
ACTGCTGAAAGTGGTGAAGTTCGGAGAGGTCATCAGCTACAGCCACCTGGCCG
CCCTGGCCGGCAATCCCGCCGCCACCGCCGCCGTGAAAACCGCCCTGAGCGG
AAATCCCGTGCCCATTCTGATCCCATGTCATCGAGTAGTGCAAGGAGATCTGGA
CGTGGGGGGCTACGAGGGCGGGCTCGCCGTGAAAGAGTGGCTGCTGGCCCAC
GAGGGCCACAGACTGGGCAAGCCTGGGCTGGGTGGCAAGCCCATCCCCAACC
CCCTGCTGGGCCTGGACAGCACCGGCGGTAAACCAATTCCAAATC-3’

The ARF1-LAP-Halo-PolyA-G418$^r$ HR plasmid was generated by linearizing the ARF1-Halo-PolyA-G418$^r$ HR plasmid with BamHI. The LAP-Halo tag was synthesized as a double strand DNA fragment and assembled into the BamHI linearized ARF1-Halo-PolyA-G418$^R$ HR plasmid via Gibson assembly.

Sequence of the double strand DNA fragment inserted:

LHA-GS-BamHI-LAP-Halo: 5’-
CTACGAAACCAGAAGGGATCGTCAGGTCGGGATCCAATGTACCCAACTTTCTTGTACAA
AGTGGTACCGGTCGCCACCGCAAGCGAGAATCTTTATTTTCAGGGCGCCGCCAAATTCA
AAGAAACCGCTGCTGCTAAATTCGAACGCCAGCACATGGACAGCGGAGGTGGAGGTTC
AAGTGGTCCCTCAGGTTCGTCGAGCCTGGAAGTTCTGTTCCAGGGGCCCCTGTCGTCT
AGCGGTCCCTCGGGTTCTATGGGCTCAGGTTCTGGAATGGCAGAAATCGGTACTGGCT
TTC-3’

To generate the SNAP-CLC HR plasmid, SNAP-Sec61Beta (Bottanelli et al., 2016) was digested with BglII and MluI and the right homology arm (~ 1Kb) was synthesized by IDT technologies as a double strand DNA fragment and assembled into the linearized vector. The resulting vector was then digested with AseI and NheI and the left homology arm (~ 1Kb) was assembled using the same strategy. Similar to previously generated constructs, the Puromycin$^r$ resistance cassette flanked by LoxP sites was introduced into the SNAP-CLC HR plasmids as a NheI-NheI fragment. The LoxP-Puromycin$^r$-LoxP fragment was amplified from the pPUR vector with the following oligoes: NheI-LoxP-Puro$^r$-sense 5’-ATGTCGCTAGCATAACTTCGTATAGCATACATTATACGAAGTTATCTGTGGAATG
TGTGTCAGTTAGGGTGTGGAAAGTCCCCAGGCTCC-3’ and Puro$^r$-LoxP-NheI-anti-sense 5’-TGCACGCTAGCATAACTTCGTATAATGTATGCTATACGAAGTTATCAGACATGAT
AAGATACATTGATGAGTTTGGACAAACCACAAC-3’.

To generate the BetaCOP-SNAP HR plasmid, the sequence containing the left and right homology arms (ca. 1 Kb each) to target the COPB1 locus were synthesized by Thermo Fisher with NheI and BamHI sites to insert the desired tag and a glycin/serin rich linker (GSGSGSGSGS). The SNAP-tag was cloned into the linearized plasmid as a NheI-BamHI fragment as previously described for the ARFs-SNAP HR plasmids. To select for positive clones with G418, the G418 resistance cassette was introduced as a BamHI-PolyA-G418$^r$-BamHI fragment. The resistance cassette was amplified from the ARF1-Halo HR plasmid using the following primers: BamHI-PolyA-sense 5'-ATGTCGGATCCCCGCGACTCTAGATCATAATCAGC-3' and G418$^r$-BamHI-anti-sense 5'-TCATGGGATCCTTTATTCTGTCTTTTTATTGCCGTC-3'.

HR plasmids and their respective guide RNAs plasmids were introduced into HeLa CCL-2 cells using FuGENE (Promega). G418 (1 mg/mL) and Puromycin (1 µg/mL) were added to the cells 3 days after transfection. After selection and recovery cells were either prepared for downstream applications or they were again transfected with 2 µg of pBS598 EF1alpha-EGFPcre plasmid (Addgene plasmid #11923) encoding for cre recombinase to excise the resistance cassette, in case a resistance cassette flanked by LoxP was used.

To obtain single cell clones of the ARF3- ARF4 and ARF5-Halo KIs, cells were subjected to fluorescent-activated cell sorting (FACS) and single cells were deposited in a well of a 96-well plate. Clones were genotyped via western blot using anti-Halo and anti-ARF antibodies, fluorescent microscopy and PCR using the following oligoes: ARF3-genotyping-sense 5'-GGTACATTCAGGCCACCTGTGCCACCAGCGGGGACGGG-3', ARF3-genotyping-anti-sense 5'-CTCCCACCTTCTTCCCTTAATCTCACCAACACGGAAGGGGC-3', ARF4-sense 5'-TGTGAGATGCAAGAGGTGTG-3', ARF4-genotyping-anti-sense 5'-CCAGCCAGAGAAAGATCCAAAACAC-3', ARF5-sense 5'-TTCCCCACAGTGGTATGTCCA-3' and ARF5-genotyping-anti-sense 5'-ATACCCTCAGCTCAGCTTCTCCCTC-3'. In case of ARF3, the genotyping PCR did not yield results and the selection of the ARF3 clone (clone 19) for the experiments was based on the fluorescent intensity taken out of the microscopy images (brightest clones) In case of ARF4 and ARF5, homozygous clones could be selected, based on the genotyping PCR results (ARF4: clone 7 and ARF5: clone 14).

For the generation of KI cell lines in HAP1 cells, the ARF1/3/4/5-Halo HR plasmid and their respective guide RNAs plasmids were transfected using a NEPA21 electroporation system (3 x $10^6$ cells, mixed with 5 µg of guide RNA plasmid and 5 µg of HR plasmid). Again, G418 (3 mg/mL) was added 3 days after transfection. After selection and recovery ARF3/4/5-Halo edited cells they were electroporated with 10 µg of pBS598 EF1alpha-EGFPcre to excise the resistance cassette. ARFs-Halo HAP1 cells were also sorted based on cell size and fluorescence to obtain an homogenous population of edited haploid cells (Beigl et al., 2020).

The ARF1$^{EN}$-Halo and BetaCOP$^{EN}$-SNAP double KI cell line was generated in Bottanelli et al., 2017. ARF3/ARF4/ARF5-Halo and BetaCOP-SNAP double KI were generated by transfecting BetaCOP-SNAP-polyA-G418 and BetaCOP guide in the sorted ARFs-Halo KIs.

The ARF1/3/4/5$^{EN-}$Halo and SNAP-CLCa$^{EN}$ double KI cell lines were generated by transfecting LoxP-Puromycin$^{r}$-LoxP-SNAP-CLCa HR plasmid and CLCa guide into the sorted ARFs-Halo KIs. Similarly, the ARF3$^{EN}$-LAP-Halo and SNAP-CLCa$^{EN}$ double KI cell line was generated by transfecting the CLCa HR and guide plasmids mentioned above into the ARF3$^{EN}$-LAP-Halo cell line.

To generate the ARF1-SNAP and ARF4-Halo double KIs, ARF4-Halo KI was transfected with the ARF1-SNAP-LoxP-G418$^{R}$-LoxP plasmid and the ARF1 guide. The ARF1$^{EN}$-LAP-Halo and ARF3$^{EN}$-LAP-SNAP double KI cell line was generated by transfecting the respective ARF3 reagents into the ARF1$^{EN}$-LAP-Halo cell line.

The ARF1/ARF3/ARF4/ARF5-2xALFA cell lines were generated by transfecting the HR plasmids and guides into HeLa cells.

To generate the ARF4-2xALFA and ARF1-2xV5 double KI cell line, the ARF4-2xALFA-loxP-Puro-LoxP HR plasmid was transfected with the ARF4 guide into an already existing ARF1-2xV5 cell line. Similarly, the ARF4-2xV5 and ARF5-2xALFA double KI cell line was generated by transfecting the ARF4-2xV5-loxP-Puro-LoxP HR plasmid with the guide into the already existing ARF5-2xALFA cell line.

All the cell lines generated in this study were validated via Western Blot analysis and PCR amplification and sequencing of the edited locus after antibiotic selection and/or excision of the resistance cassette.

**Table 1. List of dyes used for live cell imaging**

| Halo substrates | | | SNAP substrates | | |
|---|---|---|---|---|---|
| **Name** | **Manufacturer** | **Reference number** | **Name** | **Manufacturer** | **Reference number** |
| JF646-CA | Promega | GA1120 | SNAP-Cell 647-SiR | NEB | S9102S |
| JF571 | Lavis Lab | (Grimm et al., 2020) | JFX650 | Lavis Lab | (Grimm et al., 2021) |
| JFX650 | Lavis Lab | (Grimm et al., 2021) | JF585 | Lavis Lab | (Grimm et al., 2017) |
| JF503 | Lavis Lab | (Grimm et al., 2020) | | | |
| JF552 | Lavis Lab | (Zheng et al., 2019) | | | |

**Table 2. List of antibodies used in this study**

| Primary | Origin | Manufacturer | Reference number |
|---|---|---|---|
| anti-ALFA | mouse | NanoTag | N1582 |
| anti-ALFA | rabbit | NanoTag | N1583 |
| anti-V5 | rabbit | Cell signaling | D3H8Q |
| anti-Golgin97 | rabbit | Abcam | ab84340 |
| anti-GM130 | mouse | BD transduction laboratories | 610823 |
| anti-β'COP (CM1) | mouse | Gift from Rothman Lab | (Palmer et al., 1993) |
| anti-ERGIC53 | rabbit | Sigma | E1031 |
| anti-CHC | mouse | Novus | NB300-613 |
| anti-CLC | rabbit | Merck | AB9884 |
| anti-GRASP65 | rabbit | Thermo | PA3-910 |
| anti-βActin | mouse | Cell signaling | 8H10D10 |
| anti-Halo | mouse | Promega | G921A |
| anti-ARF1 | Mouse | Abcam | Ab2806 |

| Secondary | Manufacturer | Reference number |
|---|---|---|
| anti-mouse ATTO647N | Sigma | 50185-1ML-F |
| anti-rabbit ATTO647N | Sigma | 50185-1ML-F |
| anti-mouse Alexa Fluor 594 | Sigma | BCBJ2142V |
| anti-rabbit Alexa Fluor 594 | Sigma | 77671-1ML-F |
| anti-mouse HRP | Santa Cruz | SC-2005 |
| anti-mouse Alexa Fluor 488 | Invitrogen | A21202 |
| anti-rabbit Alexa Fluor 488 | Invitrogen | A11070 |
| anti-mouse Alexa Fluor 568 | Invitrogen | A11031 |
| anti-rabbit Alexa Fluor 568 | Invitrogen | A11036 |
